# Cancer therapy via neoepitope-specific monoclonal antibody cocktails

**DOI:** 10.1101/2024.08.02.606410

**Authors:** Colin J. Hartman, Asmaa O. Mohamed, Girja S. Shukla, Stephanie C. Pero, Yu-Jing Sun, Roberto S. Rodríguez, Nicholas F. Genovese, Nico M. Kohler, Thomas R. Hemphill, Yina H. Huang, David N. Krag, Margaret E. Ackerman

## Abstract

**Background:** Cellular heterogeneity presents a significant challenge to cancer treatment. Antibody therapies targeting individual tumor-associated antigens can be extremely effective but are not suited for all patients and often fail against tumors with heterogeneous expression as tumor cells with low or no antigen expression escape targeting and develop resistance. Simultaneously targeting multiple tumor-specific proteins with multiple antibodies has the potential to overcome this barrier and improve efficacy, but relatively few widely expressed cancer-specific antigens are known. In contrast, neoepitopes, which arise from mutations unique to tumor cells, are considerably more abundant. However, since neoepitopes are not commonly shared between individuals, a patient-customized approach is necessary and motivates efforts to develop an efficient means to identify suitable target mutations and isolate neoepitope-specific monoclonal antibodies.

**Methods:** Here, focusing on the latter goal, we use directed evolution in yeast and phage display systems to engineer antibodies from non-immune, human antibody fragment libraries that are specific for neoepitopes previously reported in the B16F10 melanoma model.

**Results:** We demonstrate proof-of-concept for a pipeline that supports rapid isolation and functional enhancement of multiple neoepitope peptide-targeted monoclonal antibodies and demonstrate their robust binding to B16F10 cells and potent effector functions *in vitro*. These antibodies were combined and evaluated *in vivo* for anti-cancer activity in tumor-bearing mice, where they suppressed B16F10 tumor growth and prolonged survival.

**Conclusions:** These findings emphasize the potential for clinical application of patient-customized antibody cocktails in the treatment of the many cancers poorly addressed by current therapies.

## INTRODUCTION

Current cancer antibody therapies often focus on targeting a single overexpressed cell-surface antigen; however, cancer cell populations are heterogeneous and cells with low or no expression of the target antigen do not bind to quantities of antibody sufficient for clearance by host white blood cells [1]. This phenomenon, known as antigenic escape, allows these cells to continue proliferating, ultimately undermining the efficacy of the treatment [1–3]. Epidermal growth factor receptor (EGFR) and human epidermal growth factor receptor 2 (HER2) are examples of common targets for antibody therapies that often fail to provide a lasting response [1, 4, 5] and fail to offer high cure rates [1]. Furthermore, current antibody therapies can only be used on a small subset of eligible patients with an overexpressed antigen, which may also be expressed on healthy tissues, spurring concerns of on-target off-tumor toxicity [6].

Cancer neoantigens, which are altered proteins generated by non-synonymous mutations originating in tumor cells [7], are abundant in multiple tumor types but overlooked in the design of antibody therapeutics since they are rarely shared between individuals, are not driver mutations that provide a growth advantage, and are often not overexpressed [1, 8–11]. However, neoantigens offer ideal immunological targets since they can be more immunogenic than tumor associated antigens, are not expressed on healthy cells, and may be less likely to induce autoimmunity [6, 12]. Simultaneously targeting multiple cancer neoantigens has the potential to overcome tumor heterogeneity by facilitating complete tumor coverage with antibodies, leading to robust tumor clearance [1, 9, 13, 14]. Moreover, antibody combinations may have potential additive or synergistic effects [14, 15].

Cancer vaccines and adoptive cell transfer approaches targeting neoantigens have been explored [6, 16–21], however, an antibody approach has yet to be established in the clinic. Patient-customized cancer antibody therapy was first investigated in the early 1980s in the form of anti-idiotype antibodies for the treatment of B-cell lymphoma [2, 22]; however, the laborious amount of work and cost associated with producing customized antibody therapies led to investigation of targeting more generic B cell markers, and the safety and success of anti-CD19 and anti-CD20 antibodies reduced enthusiasm for further exploring this approach [23].

Nonetheless, we have previously shown that a cocktail of polyclonal antibodies specific for melanoma or breast cancer cell surface mutations improves the survival of tumor-bearing mice [1, 24]. This study strives to recapitulate the success of the cocktails of polyclonal rabbit-derived anti-sera with monoclonal antibodies by adapting efficient library display and affinity maturation methods to engineer and then functionally optimize humanized antibodies.

We report use of yeast surface display (YSD) and phage display (PD) technologies to identify antibodies targeting accessible neoepitopes in B16F10 melanoma cells. By simultaneously targeting multiple neoantigens, we aim to develop a therapeutic approach that can overcome tumor heterogeneity and reduce the likelihood of antigenic escape. This study also explores the potential for combining these neoantigen-targeting antibodies with immune checkpoint blockade (ICB) therapies to enhance therapeutic efficacy. Ultimately, we aim to build a pipeline for engineering robust, multi-neoepitope targeted antibody cocktails that can contribute to more effective and durable cancer treatment, paving the way for clinical translation and improved patient outcomes.

## RESULTS

### Yeast surface display for neoepitope-targeted antibody discovery

We utilized a naïve or nonimmune human single-chain variable fragment (scFv)- expressing YSD library (diversity ≈ 1x10^9^) [25] with the aim to discover scFvs targeting nine previously described mutated peptide sequences identified from whole exome sequencing of the B16F10 cell line [1, 24, 26] (**Fig. 1A-B**). These neoepitope peptides were synthetized in the context of 11-mer peptides, with the mutation typically located at the center of the peptide and native amino acids flanking the mutation. Using gene ontology analyses from UniProt and QuickGO, some of the targeted proteins are membrane bound while others are secreted into the extracellular space but remain associated with the cell membrane. Using a series of positive and negative selections accomplished by magnetic bead and flow cytometry-based selections and one round of mutagenesis (**Fig. 1C**), clones specific for four of the nine candidate neoepitope peptides were isolated (**Fig. 1D**). Positive selections used biotinylated target peptides bound to streptavidin (SA) beads or tetramerized peptides. Negative selections against SA-only beads and off-target peptides were conducted to prevent reagent and non-specific binding. One round of mutagenesis via error-prone PCR was conducted for each target peptide. Multiple clones from each enriched library were considered for downstream functional analyses. These candidates were cloned into mammalian expression vectors as fully human scFv-Fc IgG1 antibodies (**Fig. 1A**).

**Figure 1.**
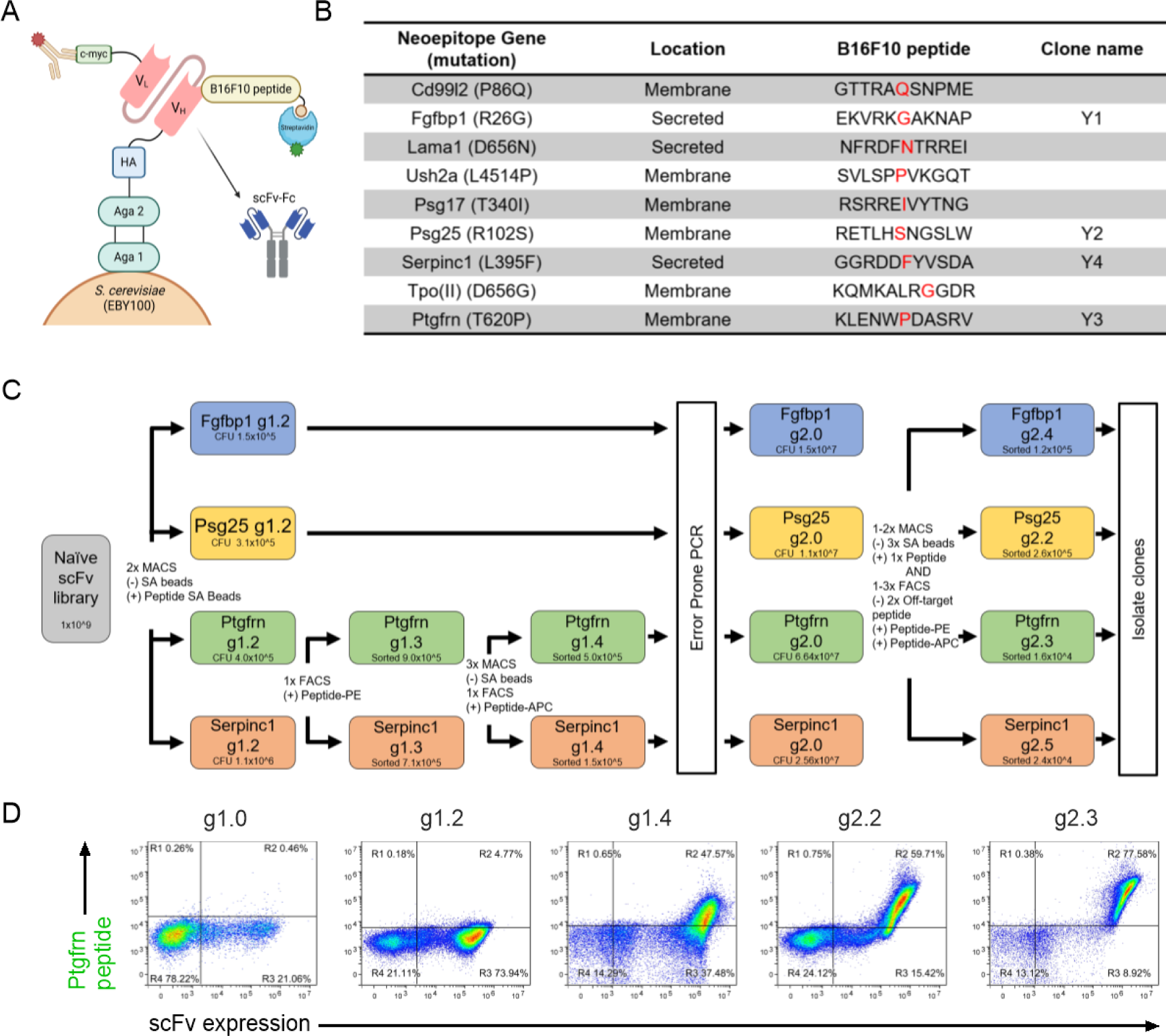
Overview of the single-chain variable fragment (scFv) yeast display library screening pipeline. **A.** Graphical schematic showing the nonimmune human scFv-expressing yeast display library format, B16F10 neoepitope peptide and c-myc tag expression staining strategies, and reformatting of selected clones as scFv-Fc fusion proteins. **B.** List of neoepitope 11-mer peptides screened against the yeast display library and resulting clones (Y1-Y4). **C.** Process diagram showing the sequential steps in the isolation of peptide-specific scFvs from the yeast display library for each of four neoepitope peptides. Population generations (g) and sizes indicated in boxes for each peptide. Positive (+) and negative (-) selection and diversification processes indicated between boxes. **D.** Exemplary flow cytometry biplots of scFv expression versus peptide (Ptgfrn) binding over the course of serial selections for indicated populations. V_L_, variable light chain; V_H_, variable heavy chain; HA, hemagglutinin; MACS, magnetic-activated cell sorting; FACS, fluorescence-activated cell sorting; CFU, colony forming units; SA, streptavidin.

### Yeast display antibody validation

Candidate clones from each enrichment pool were screened for binding to both mutant and wild-type peptides, as well as for binding to B16F10 cells. Although most candidate antibodies bound well to the neoepitope peptides, some clones did not bind effectively to the target B16F10 cells (data not shown), potentially due to conformational differences between short-synthetic peptides and expression within the context of the entire protein and cell. For each peptide, the antibody clone with the most favorable properties (i.e., high mutant peptide binding, low wild-type peptide binding, and binding to B16F10 cells) was chosen for downstream use (Y1 - Fgfbp1, Y2 - Psg25, Y3 - Ptgfrn, and Y4 - Serpinc1).

Flow cytometry was used to determine binding to mutant and wild-type peptides. All clones bound to the mutant peptides on SA beads; however, most clones also showed some level of binding to the wild-type peptides (**Fig. 2A**). Fluorescence microscopy indicated that each antibody independently and as a four-antibody cocktail bound to B16F10 cells (**Fig. 2B-C**).

**Figure 2.**
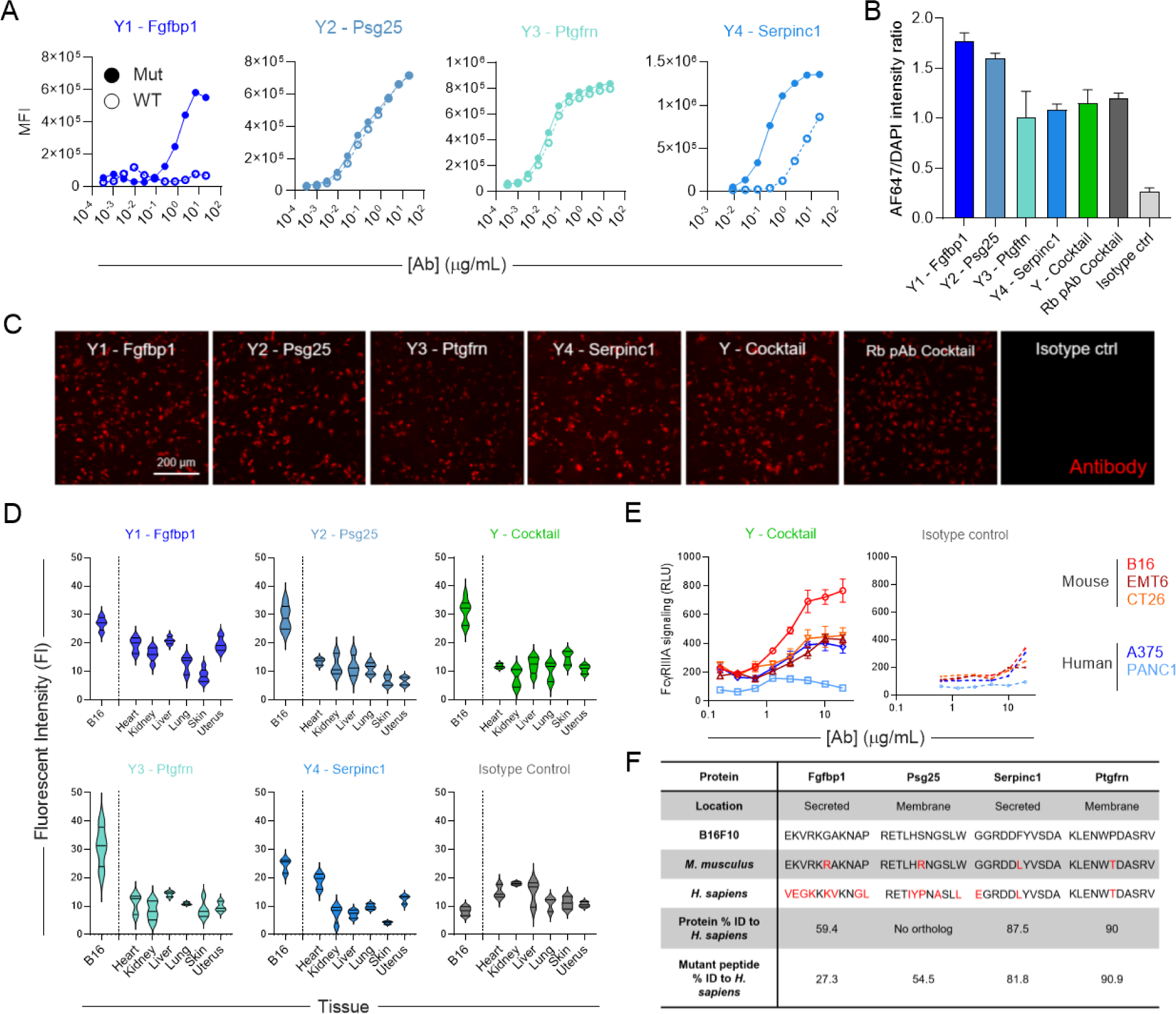
Neoepitope-targeted yeast-derived antibodies bind to B16F10 mutated peptides and cells. **A.** Binding of yeast-derived antibodies (Y1-Y4) to wildtype (hollow) and mutant (filled) peptides. **B-C.** Quantification of fluorescent microscopy staining of B16F10 cell line with indicated antibodies at 10 µg/mL (**B**), and representative fluorescent microscopy staining images of B16F10 cells (**C**). **D.** Binding of yeast-derived antibodies to malignant B16F10 tumor and a panel of normal tissues from C57B/6 mice. Data are expressed as median with a 95% CI of 3-6 replicates. **E.** Antibody-induced FcγRIIIa signaling measured using Jurkat-Lucia NFAT- CD16 reporter cells for target human (red) and mouse (blue) cell lines. **F.** Analysis of peptide and protein conservation between B16F10, wildtype *M. musculus*, and *H. sapiens* orthologs/proteome. Amino acids in red indicate differences from B16F10. Unless otherwise noted, error bars represent SD of 3-5 replicates. WT, wild type; MFI, median fluorescence intensity; Rb pAb, positive control rabbit polyclonal antibody cocktail, RLU, relative light units; ID, identity.

The rabbit polyclonal antibody (Rb pAb) cocktail, which targets the same nine mutated B16F10 peptides, has been demonstrated to bind to B16F10 cells, and has shown efficacy in B16- bearing mice [1, 24] served as a positive control, while a randomly selected scFv-Fc was employed as a negative control.

Since the mutant and wild-type peptides differ by only one amino acid and most antibodies bound to both peptides, we explored off-target binding of these antibodies to a panel of wild-type mouse tissues including heart, kidney, liver, lung, skin, and uterus. While each antibody exhibited the highest binding to B16F10 tumor sections, some off-target tissue binding was observed (**Fig. 2D, Supplementary Fig. 1**), motivating use of a cocktail to increase binding to tumor relative to normal tissue. Interestingly, this off-target binding decreased when using an equivalent mass quantity of the antibody cocktail, suggesting that a cocktail may help mitigate off-target effects. Although any single target protein with the wild-type epitope could be expressed in other tissues, it is unlikely that all wild-type epitopes would be co-expressed in the same location, thereby limiting the binding sites in any one tissue. Conversely, all these targets are expected to be present at the tumor site.

We further examined off-target effects by measuring the functional activity of these antibodies as a cocktail against mouse cell lines expressing the wild-type epitopes and human cell lines that may express proteins with similar sequences in the form of an FcγRIIIa signaling assay using Jurkat-Lucia NFAT-CD16 reporter cells [27, 28] as a proxy for antibody-dependent cellular cytotoxicity (ADCC) (**Fig. 2E-F**). As compared to other cell lines, the antibody cocktail induced the highest CD16-dependent FcγRIIIa signaling against B16F10 cells. Taken together, these data demonstrate that B16-targeted YSD-derived antibodies bind to B16F10 cells with limited off-target binding.

### Antibody engineering for enhanced effector functions

Individually, all but one of the B16-targeted YSD-derived antibodies exhibited CD16- dependent FcγRIIIa signaling activity exceeding even the rabbit polyclonal antibody cocktail (**Fig. 3A**). When combined into a cocktail, while maintaining the same total mass of antibody, ADCC activity improved. A synergistic effect at concentrations of 1.25 µg/mL and above was demonstrated using the Bliss Independence Model [29].

**Figure 3.**
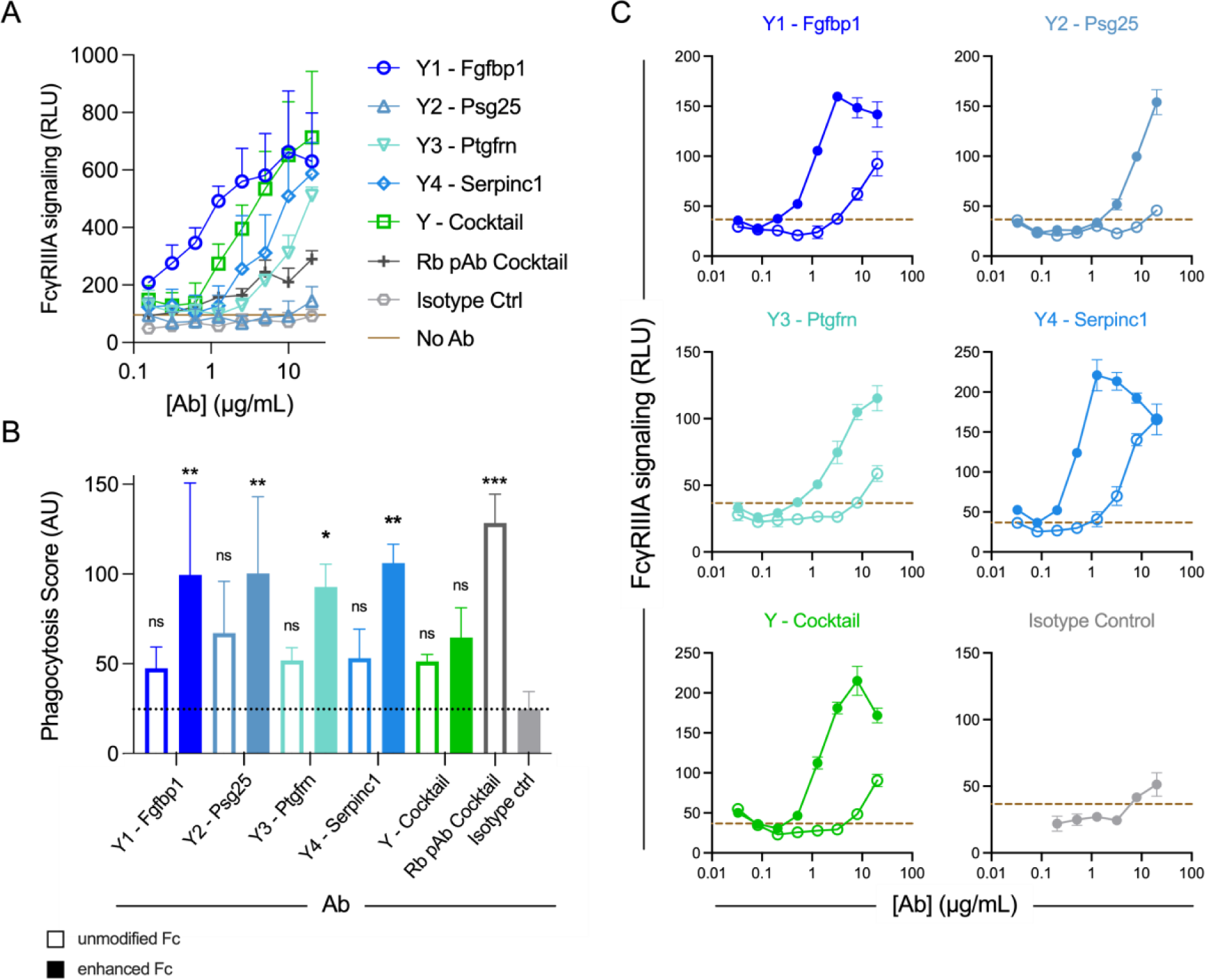
Neoepitope-targeted yeast-derived antibodies elicit effector functions. **A.** Antibody-induced FcγRIIIa signaling measured using Jurkat-Lucia NFAT-CD16 reporter cells for target B16F10 cells. **B.** Antibody-dependent phagocytosis of B16F10 cells by THP-1 monocytic cell line for unmodified (hollow bar) and Fc enhanced (filled bar) antibodies. The horizontal dotted line represents the mean of the isotype control. A one-way ANOVA (*F*_11,24_ = 5.117, *p* = 0.0004) followed by Dunnett’s post hoc test was used to compare groups to the Isotype control (ns *p* ≥ 0.05, * *p* < 0.05, ** *p* < 0.01, *** *p* < 0.001). **C.** Comparison of FcγRIIIa signaling activity when Fc-enhanced (filled) and unmodified (hollow) antibodies were co-cultured with B16F10 cells. Error bars represent SD of 3 replicates. Rb pAb, positive control rabbit polyclonal antibody cocktail; RLU, relative light units; AU, arbitrary units.

Phagocytosis of B16F10 cells by THP-1 monocytes was monitored by flow cytometry (**Supplementary Fig. 2A**). Notably, complete engulfment of B16F10 cells was not observed; rather, trogocytosis appeared to be the method of phagocytic activity of the THP-1 cells, potentially due to their small cellular diameter (10.6 ± 1.5 µm) [30] as compared to larger B16F10 cells (15.4 ± 1.4 µm) [31] (**Supplementary Fig. 2B**). Whereas the rabbit polyclonal cocktail exhibited robust phagocytic activity, YSD-derived antibody phagocytic activity was not significantly elevated over that of the isotype control (**Fig. 3B**), motivating the use of antibody engineering strategies to increase effector function.

To enhance the potency of these B16-targeted YSD-derived antibodies, we engineered their Fc regions to contain the GASDALIE (Gly236Ala/Ser239Asp/Ala330Leu/Ile332Glu) mutation set which is known to enhance ADCC and antibody-dependent cellular phagocytosis (ADCP) activity by increasing binding to FcγRIIIa and FcγRIIa [32–36]. Fc-enhanced antibodies, denoted by a subscript “E,” each demonstrated higher ADCC and ADCP activities compared to their wild-type Fc counterparts and negative controls (**Fig. 3B-C**). Notably, whereas the wild-type Fc antibody Y2 - Psg25 had no ADCC activity, after Fc engineering, it showed ADCC activity at high concentrations. In most cases, phagocytic activity increased three-fold, while ADCC activity increased by an order of magnitude for antibodies with Fc-enhancing mutations.

### Phage surface display for neoepitope-targeted antibody discovery

In parallel, antibody fragments were also isolated from a nonimmune, human, Fab PD library (diversity ≈ 1x10^11^) (**Fig. 4A**) to engineer antibodies targeting four of the nine B16F10 neoepitope peptides. A set of four peptides (**Fig. 4B**) were selected on the basis of high tumor binding with Rb pAbs [1, 24], over three rounds of panning (**Fig. 4C**). Selections were conducted using unmodified and variably conjugated forms of the peptides with the aims of diversifying the modes of peptide presentation and maximizing chances of success. One phage-derived (P) antibody candidate for each target was ultimately chosen for downstream use and cloned into mammalian expression vectors as fully human IgG1 kappa antibodies with and without the GASDALIE mutation set, denoted by a subscript “E,” to enhance their effector function.

**Figure 4.**
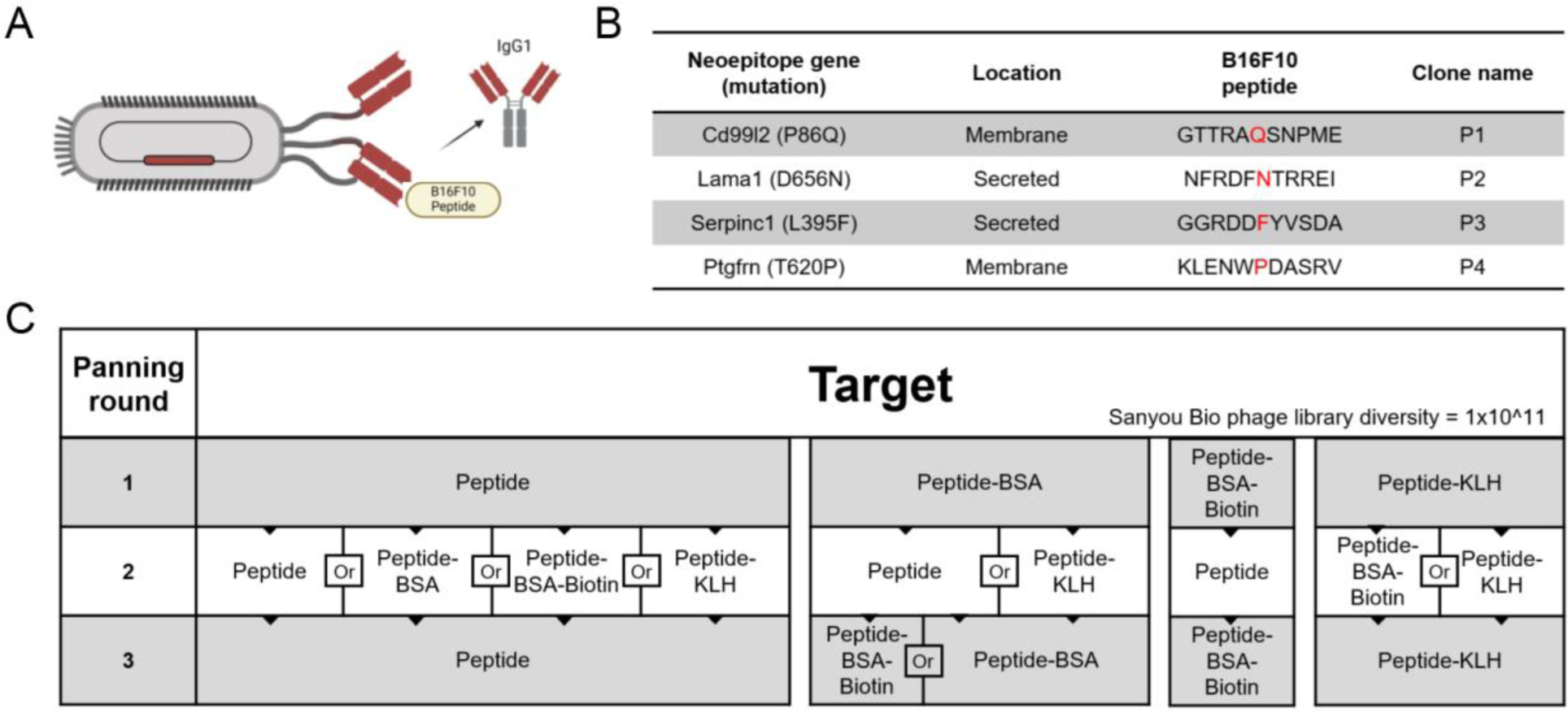
Overview of the Fab phage display library screening pipeline. **A.** Graphical schematic showing the nonimmune, human, Fab-expressing phage display library format and reformatting of selected clones as antibodies. **B.** List of four neoepitope 11-mer peptides screened against the phage display library and resulting clones (P1-P4). **C.** Process flowchart showing the sequential steps in the isolation of Fab fragments specific for each peptide from the phage display library. BSA, bovine serum albumin; KLH, keyhole limpet hemocyanin.

### Phage display antibody validation

Each of the selected antibodies exhibited binding to one or more forms of the mutant peptides either in multiplexed bead assays (**Fig. 5A**) or in ELISAs (**Fig. 5B**). Notably, some antibodies (P2_E_ - Lama1 and P4_E_ - Ptgfrn) showed no detectable binding to either the mutant or wild-type peptides in the absence of conjugation. Nonetheless, all antibodies were also shown to bind to B16F10 cells using flow cytometry (**Fig. 5C**) and immunocytochemistry (**Fig. 5D**).

**Figure 5.**
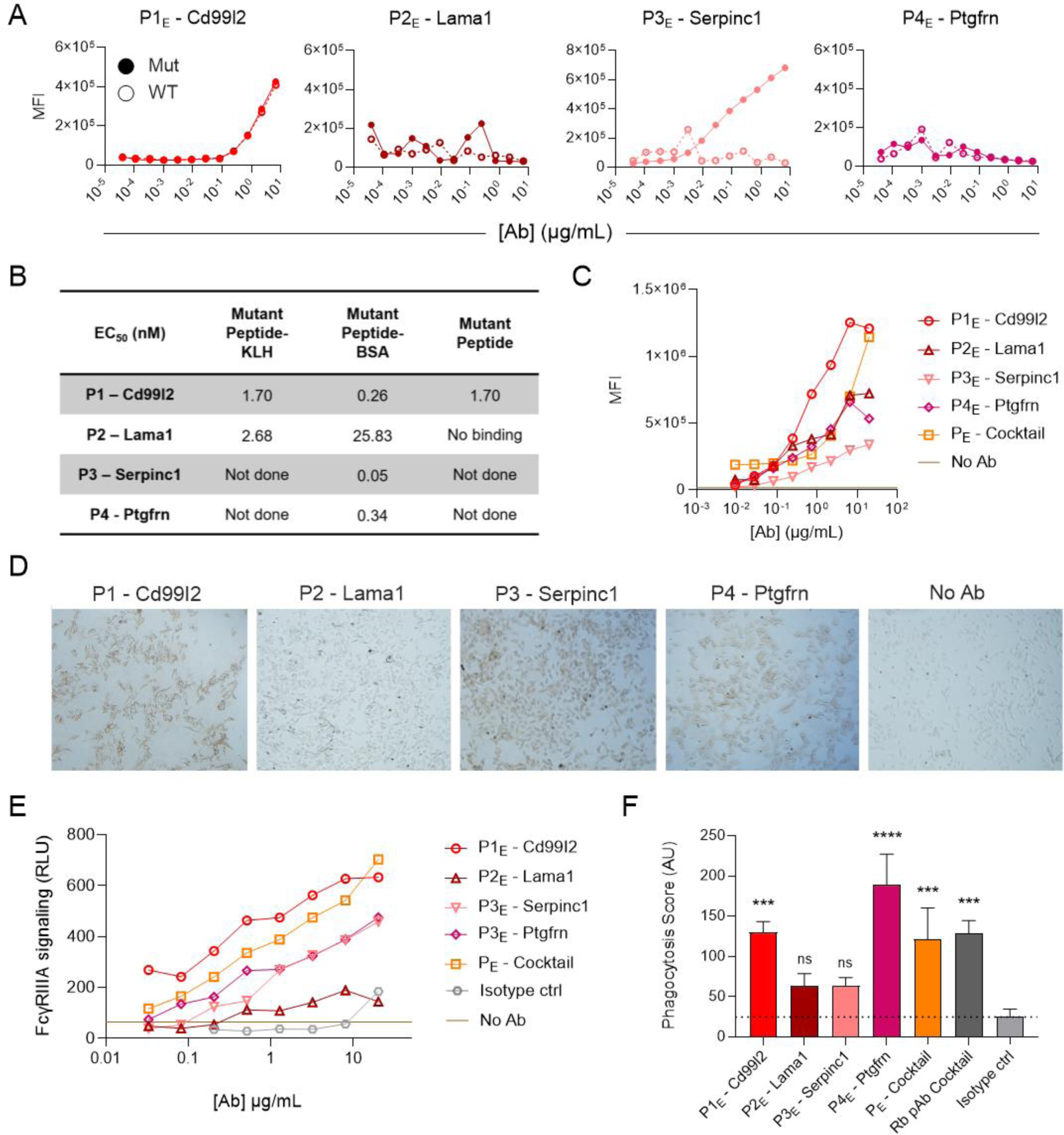
Neoepitope-targeted phage-derived antibodies bind to B16F10 mutated peptides and cells and elicit effector functions. **A-B.** Binding of phage-derived (P) antibodies (P1-P4) to wildtype (hollow) and mutant (filled) peptides by multiplex assay (**A**) and ELISA (**B**)**. C-D.** Flow cytometric (**C**) and immunohistochemistry (**D**) staining of B16F10 cells. **E.** Antibody-dependent FcγRIIIa signaling measured using Jurkat-Lucia NFAT-CD16 reporter cells. **F.** Antibody-dependent phagocytosis of B16F10 cells with THP-1 monocytes. A one-way ANOVA (*F*_6,14_ = 16.78, *p* < 0.0001) followed by Dunnett’s post hoc test was used to compare groups against the isotype control (ns *p* ≥ 0.05, * *p* < 0.05, ** *p* < 0.01, *** *p* < 0.001, **** *p* < 0.0001). Error bars represent SD of 3 replicates. Phage-derived antibodies with unmodified Fc domains are indicated as P and Fc enhanced Fc domains as P_E_. MFI, median fluorescence intensity; EC_50_, half-maximal effective concentration, BSA, bovine serum albumin; KLH, keyhole limpet hemocyanin; Rb pAb, positive control rabbit polyclonal antibody cocktail.

Additionally, the enhanced versions of these PD-derived antibodies elicited ADCC activity in the form of FcγRIIIa signaling (**Fig. 5E**) and phagocytic activity (**Fig. 5F**). Together, these data show that the phage-derived antibodies bound to B16F10 cells and effectively elicited multiple antibody effector functions supporting their investigation *in vivo*.

### Inhibiting tumor growth with antibody cocktails

In initial *in vivo* experiments, B16F10-bearing C57BL/6 mice were treated with the PD-derived antibody cocktail lacking Fc enhancing mutations in combination with anti-mouse PD-1 ICB (**Supplementary Fig. 3A**). At the end of the experiment, 4, 1, 1, and no mice survived in the cocktail, isotype control, ICB alone, and PBS-treated control groups, respectively (**Supplementary Fig. 3B**). Tumor volume increased most among PBS-treated animals, was similar in ICB alone and isotype control groups, and was lowest in the cocktail-treated group (**Supplementary Fig. 3C-D**).

Similarly, an initial experiment with the YSD-derived antibody cocktail lacking Fc-enhancing mutations was conducted in which both low (50 µg per antibody) and high (200 µg per antibody) doses of the cocktail, or with a single antibody (Y4 – Serpinc1, 200 µg) were combined with anti-mouse PD-1 ICB (**Supplementary Fig. 4A**). As compared to buffer and isotype controls, a greater number of animals survived to the predetermined endpoint of the experiment (**Supplementary Fig. 4B**). Additionally, tumor growth was also slower in animals treated with the YSD-derived antibody cocktails or clone, although this was not significant (**Supplementary Fig. 4C**-D). In this experiment, surviving mice were sacrificed and their tumors excised and weighed on day 20 post tumor implantation, showing a decrease in tumor weight in the anti-B16F10 antibody treated groups (**Supplementary Fig. 4E**).

A follow-up experiment that also included the Fc-enhanced YSD-derived cocktail was conducted to evaluate the effect of these Fc-enhancing mutations. Mice were treated similarly to the previous regimen but with the addition of an anti-mouse CTLA-4 ICB antibody in combination with anti-mouse PD-1 (**Supplementary Fig. 5A**). Although some mice survived in both the wild-type Fc and Fc-enhanced antibody cocktail groups, whereas none survived in the IgG control group, this difference was not statistically significant (**Supplementary Fig. 5B**), and tumor growth kinetics were similar in all groups treated with ICB (**Supplementary Fig. 5C-D**).

Intriguingly, one surviving mouse in the wild-type Fc cocktail group presented with vitiligo at the site of tumor implantation around day 70 (**Supplementary Fig. 5E**), suggesting the possibility of a vaccinal effect. Vitiligo is characterized by the loss of melanocytes and can indicate a memory T cell response [37–39]. To investigate this possibility, the surviving mice were re-challenged with B16F10 cells but not retreated with antibodies 117 days after the initial tumor implantation.

These mice exhibited delayed tumor growth compared to a control group of untreated mice (**Supplementary Fig. 5F**). Indeed, no tumor growth was observed in the animal treated with the enhanced cocktail originally. These data suggest that epitope spreading occurred, generating a robust immune response not only against the targeted neoepitopes but also against other melanoma or melanocyte epitopes.

Encouraged by these preliminary results, we aimed to improve study design and increase statistical power. In this new study, mice were implanted with B16F10 cells and treated every other day for a total of five treatments starting on day 5 with either an IgG antibody control, the Fc-enhanced YSD-derived antibody cocktail, or the Fc-enhanced PD-derived antibody cocktail (**Fig. 6A**). On the second treatment day, a single dose of anti-mouse CTLA-4 and anti-mouse PD-1 ICB antibodies was administered. Mice receiving either YSD- or PD-derived antibody cocktails had improved survival (**Fig. 6B**) and inhibited tumor growth compared to the control group (**Fig. 6C-D**). In support of these results, this experiment was repeated and similar outcomes were observed (**Supplementary Fig. 6**). Together these data show that the strategy of targeting cancer neoepitopes with cocktails of functionally enhanced monoclonal antibodies is effective in mice, motivating further investigation of this approach.

**Figure 6.**
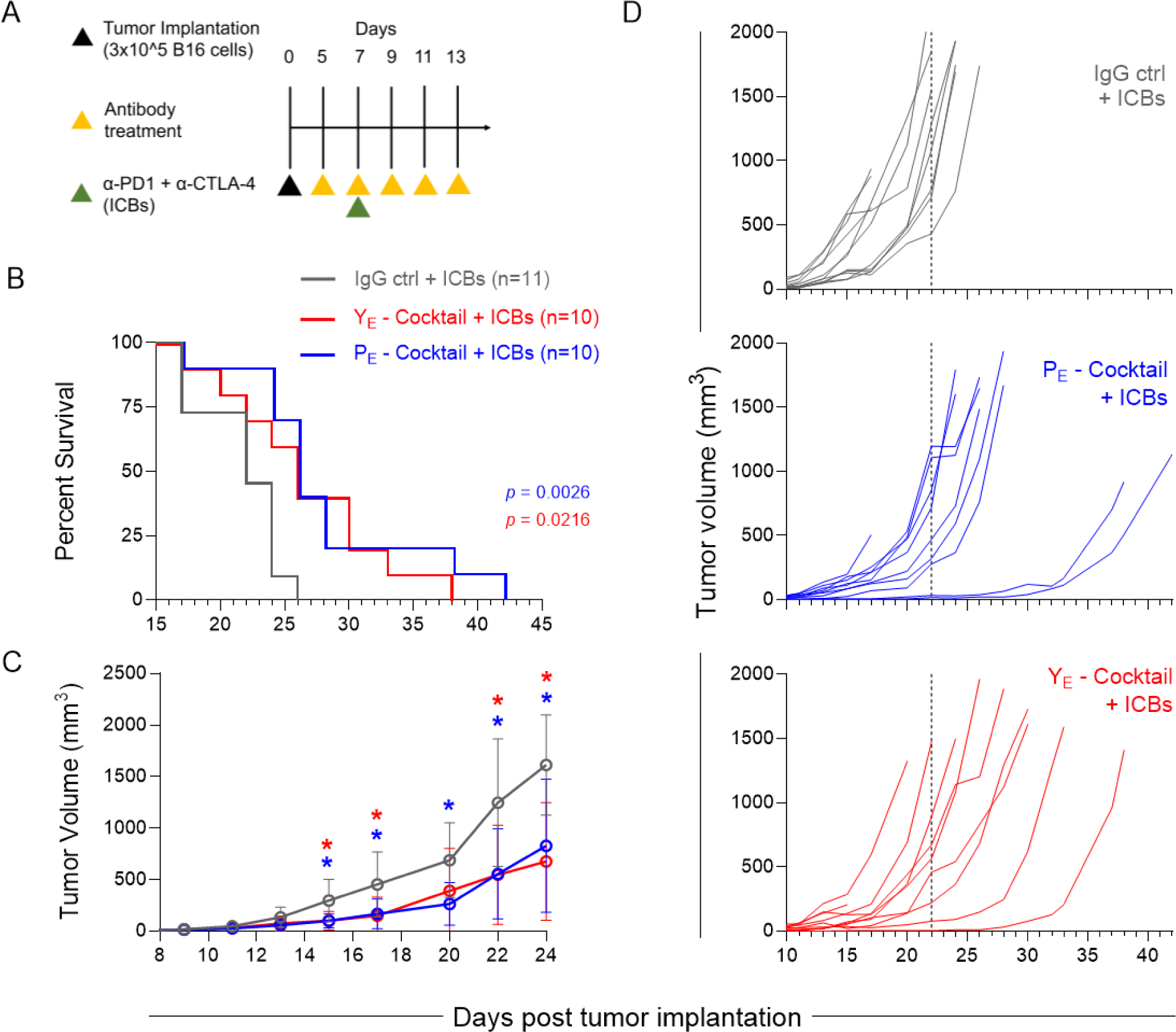
Neoepitope-targeted yeast- and phage-derived antibody cocktails inhibit tumor growth in B16F10-bearing C57BL/6J mice. **A.** Treatment regimen in which antibody cocktails or isotype control (IgG ctrl) and immune checkpoint blockade (ICBs) were administered to tumor-bearing mice. **B.** Kaplan-Meier survival curves. A log-rank Mantel-Cox test was used to calculate a difference in the survival curves for phage and yeast antibody cocktails as compared to isotype control. **C.** Tumor growth curves are depicted over time following confirmed tumor implantation. A two-way ANOVA (Time: *F*_2.1, 54.18_ = 65.82, *p* < 0.0001; Treatment: *F*_2, 28_ = 11.78, *p* = 0.0002; Time x Treatment: *F*_20, 258_ = 5.367, *p* < 0.0001) with a Greenhouse-Geisser correction (ε = 0.21) followed by Dunnett’s post hoc test was used to compare treatment groups against the IgG control + ICBs group. Error bars represent SD of the mean, \**p* < 0.05. **D.** Tumor growth curves for individual mice. The vertical line indicates the median survival of mice in the IgG control + ICBs group.

## DISCUSSION

Antibody therapies for cancer targeting one tumor antigen can be extremely effective but often fail against tumors with heterogeneous expression of tumor antigens, as tumor cells with low or no antigen expression escape targeting and develop resistance. Similarly, while immune checkpoint modulators have expanded clinical options, many patients are diagnosed with cancers for which these interventions are either not suitable or are ineffective [40–42].

Simultaneously targeting multiple tumor neoantigens unique to an individual’s tumor may overcome tumor heterogeneity, but this approach has historically been presumed to be infeasible. Perhaps ironically, the first successful monoclonal antibodies for cancer actually targeted epitopes unique to individual subjects (anti-idiotypic antibodies for B cell lymphomas) [2, 22]. However, this success has been largely forgotten in the quest to identify one-drug-for- some therapies.

Technical advancements make it realistic to revisit these early successes and attempt to generalize them across tumor types. It is now possible to sequence tumor and healthy tissues to identify cancer neoepitopes, to infer surface expression and predict the immunogenicity of mutant peptides present in putative cell surface-associated proteins, and to use robust and rapid antibody engineering platforms to develop customized cocktails of tumor-specific antibodies. While a customized approach has been previously demonstrated in the B16F10 melanoma model using polyclonal antibody pools derived from peptide-immunized rabbits [1, 24], here, as a next step toward clinical translation, we sought to explore the ability of monoclonal antibodies to recapitulate success of this approach.

Compared to immunization-based approaches for antibody development, *in vitro* methods for generating neoepitope-specific antibodies offer several potential advantages, including speed and the ability to use human libraries. While animal model-derived neoepitope-specific B cells could be isolated and cloned to produce chimeric or humanized antibodies, immunization processes usually proceed over weeks to months. Both phage and yeast platforms can support more rapid isolation of peptide-specific antibodies. Additionally, though not incorporated in the enrichment strategies employed in the proof-of-concept experiments presented here, *in vitro* methods also permit negative selections against native peptides to be conducted, which could improve the safety profile of this approach. Further, if specific affinities or other properties were found to be necessary for or associated with clinical success, *in vitro* methods permit robust affinity maturation and offer the ability to accommodate diverse design constraints.

This study demonstrated the successful isolation of neoepitope peptide-specific antibodies using both yeast and phage display methods. These antibodies targeted both B16F10 cells and tumor sections, indicating potential specificity towards mutated epitopes.

However, the degree of mutation-specificity varied, with some antibodies demonstrating cross-reactivity to native peptides. Notably, these antibodies exhibited anti-tumor activity *in vitro*, presumably through effector functions such as ADCC and ADCP, particularly when engineered with Fc function-enhancing mutations. These activities have been shown to be important for tumor clearance [43–49]. Complement-dependent cytotoxicity (CDC) is another potent antibody effector mechanism [50–52] that was not examined in this study. A better understanding of how these antibodies may regulate tumor cell killing through these mechanisms is important for holistically understanding their efficacy. Both yeast- and phage-derived antibodies showed evidence of *in vivo* activity through effective suppression of tumor growth.

However, significant limitations and challenges remain, both in this study and in the broader concept of patient-customized neoepitope-targeting antibody cocktails. These include the lack of confirmation of antigen expression in the B16F10 cells used, the lack of high antibody affinity, and variable cross reactivity to native peptides. Efficacy in animal models was inconsistent, with several studies failing to show statistically significant results, partly due to relatively low power in some models. The use of a single control antibody rather than a cocktail, and the comparison of yeast-derived scFv-Fc antibodies to a full-length IgG1 control instead of a true scFv-Fc isotype control, also presents a limitation of these findings. It is also important to recognize that the antibodies used in these *in vivo* studies are human antibodies, which likely reduces their half-life. Moreover, this treatment approach for this study was not investigated without the use of ICB nor was it examined in the context of other tumor models.

From a clinical standpoint, efficacy challenges may arise if there is an inability to confirm antigen expression or bioactivity in patient tumors beyond peptide screening. Additional uncertainties include determining the optimal number of peptide targets, which may vary between patients, the labor-intensive nature of custom antibody development for individual patients, and regulatory frameworks that are ill-suited for personalized medicine approaches.

Additionally, direct tumor injection was used for these *in vivo* models due to its potential efficacy and toxicity advantages [53–55]; however, this administration route may not always be feasible in clinical settings.

Since neoepitope mutations are rarely shared between individuals, this approach must be patient-customized. Each cancer patient would need an entirely new drug cocktail of antibodies to treat their specific tumor. Although this is a manufacturing hurdle, this approach offers unparalleled flexibility in achieving tumor-specific targeting for more patients, which could offer higher response rates and robust efficacy. This approach of targeting multiple neoepitopes may also help overcome tumor heterogeneity by reducing the probability that any one cancer cell does not express one of the target epitopes and reducing the likelihood of tumor escape.

There may also be some safety advantages, through improved therapeutic index. In this context, the therapeutic index can be considered to be proportional to the drug’s selectivity for tumor tissue over healthy tissues, which is crucial for minimizing toxicity and maximizing therapeutic efficacy: although any single target protein with the wild-type epitope could be expressed in other tissues, it is unlikely that all wild-type epitopes would be co-expressed in the same location, thereby limiting the binding sites in any one healthy tissue as compared to the tumor thereby potentially increasing the therapeutic index. Conversely, all these targets are expected to be present at the tumor site.

Furthermore, we recognize the value of incorporating a screening step to eliminate wild-type binders during the library enrichment process. While not initially integrated into these pipelines, such an addition could substantially enhance the safety and efficacy of this approach. Enhancing the speed of the pipeline through mRNA or ribosomal display could be explored to improve time to product for improved patient outcomes. To enhance the screening process of antibody candidates, cell lines expressing the target proteins could be generated or, when feasible, patient-derived target cells could be employed in development and screening. More work will be necessary to determine if Fc-enhancing mutations have added value, which dosing regimens will be most effective, and if antibody-drug conjugates could be leveraged to improve outcomes.

Exploring alternative applications of this concept to improve speed and feasibility could involve investigating mRNA nanoparticle delivery or adeno-associated vector (AAV) technologies for expression of antibodies *in vivo*. These delivery systems have shown promise in efficiently delivering genetic material for therapeutic purposes, including the expression of therapeutic antibodies [56–62]. Additionally, the utilization of predictive software tools, such as AlphaFold, represents a pivotal advancement to identify antibody-accessible neoepitopes.

These tools could help identify the most efficient targets, particularly those situated distally to the cell membrane and externally exposed residues on the protein surface. By incorporating predictive software into the screening process, it may be possible to expedite antibody discovery, enhance target specificity, and accelerate the translation of these findings into clinical applications. This approach not only saves crucial time in transitioning from laboratory research to patient treatment but also would strive to enhance efficacy, particularly given the rapid evolution and progression of many cancers [63, 64].

Alternatives also include active vaccination, such as by encoding neoantigens in patient-customized mRNA vaccines, an approach marking a significant concurrent stride in personalized cancer immunotherapy [19, 20]. Relevant to this active immunization approach, we conducted preliminary immunization experiments in mice, but failed to generate robust antibody responses or successfully clone peptide-specific antibodies, perhaps due to similarity to self, given the murine tumor targets. In contrast, recombinant antibody cocktails can potentially offer more robust efficacy. The *ex vivo* generation of antibodies ensures precise neoepitope targeting that does not depend on the variability of individual immune responses and the need to overcome potential tolerance barriers due to the nature of neoepitopes and their similarity to their wild-type counterparts. As a result, antibody cocktails may offer a compelling complementary strategy to mRNA vaccines.

In sum, both tumor and patient heterogeneity pose a significant challenge in cancer treatment. However, the development of robust and rapid antibody engineering platforms offers the prospect of revisiting and extending the success of early patient-specific antibody therapies. To this end, we report the use of two versatile antibody engineering platforms to screen monoclonal antibodies to dozens of these novel target peptides for each patient, to engineer these antibodies for enhanced anti-tumor activity *in vivo*, and to deliver them as recombinant protein or by vectored (e.g., mRNA or AAV) approaches. Here we combine and improve upon these techniques to report a pipeline that could support practical delivery of such highly efficacious antibodies to cancer patients. Through these concerted efforts, we envision the prospect of a new era of personalized cancer therapy—shifting from the current one-drug-for-many approach to a many-drugs-for-one-patient strategy—which could offer renewed hope to patients.

## METHODS

### B16F10 neoepitope peptides

Nine previously described mutated protein sequences obtained from whole exome sequencing of the B16F10 cell line [1, 24, 26] were synthesized at GenScript in the context of 11-mer peptides with N-terminal biotin and a 6-aminohexanoic acid spacer. Cellular site expression information of these mutated proteins was obtained from UniProt and QuickGO gene ontology databases. The Basic Local Alignment Search Tool (BLAST) for proteins sequences through the NIH was used to find orthologs to these mutated proteins in *Homo sapiens*.

### Isolation of neoepitope-specific antibody fragments by yeast surface display

A nonimmune human scFv-expressing yeast display library (diversity ≈ 1x10^9^) [25] in *Saccharomyces cerevisiae* was used for this work using methods similar to those previously described [65–67]. Care was taken to maintain a minimum of ten-fold the theoretical diversity of each library population. This library (generation, g1.0) utilizes galactose-inducible pCTCON2 expression vectors. Culturing was conducted in baffled flasks with shaking at 250 rpm in volumes of 15-25% of the total flask volume. Routine culture was conducted in synthetic dextrose complete amino acid (SDCAA) growth media at 30°C and cultures were induced to express scFv transgenes during log phase growth (OD_600_ = 2-5) by culturing in synthetic galactose complete amino acid (SGCAA) induction media at 20°C for 20-72 hours.

Magnetic-activated cell sorting (MACS) was conducted by capturing biotinylated peptides onto SA-coated magnetic beads (Thermo Fisher Dynabeads™ M-270). Beads were washed 5x with PBS + 0.1% BSA prior to incubation with biotinylated peptides for 2 hours at 4°C with rotation. A quantity of 500 pmol of peptide was used for every 25 µL (250 µg) of M-270 beads. After saturation with biotinylated peptides, the beads were washed 5x with PBS + 0.1% BSA to remove excess soluble peptide. An induced scFv-expressing yeast display library was used to screen for binders to each peptide. Each round of MACS consisted of one negative selection using uncoated beads for 1.5 hours and one positive selection using peptide-coated beads for 2 hours. After each round of selection, beads were washed with 1 mL of PBS + 0.1% BSA for 15 minutes to separate bead-bound yeast from yeast trapped between beads. The beads were subsequently resuspended in 1 mL of SDCAA media, 10 µL of which was used in plated serial dilutions for library diversity estimation and the remaining volume was added to 19 mL of SDCAA media + 1% penicillin-streptomycin in a 125-mL baffled flask and cultured at 30°C at 250 rpm for further selection or diversification processes.

Fluorescence-activated cell sorting (FACS) was conducted by sorting peptide tetramer-bound yeast using a Sony MA900 cell sorter. A minimum of ten-fold the theoretical diversity of the yeast library was analyzed. Tetramerized peptides were created by incubating soluble biotinylated peptides with streptavidin-PE (Southern Biotech 7105-09L) or streptavidin-APC (Southern Biotech 7105-11L) at a 4:1 molar ratio for 2 hours at 4°C with rotation. Induced yeast were stained for 1 hour with a chicken anti-c-myc eptope tag antibody (Exalpha ACMYC) and tetramerized peptides at 36 nM in PBS + 0.1% BSA. After primary incubation, cells were washed 2x with PBS + 0.1% BSA and stained with goat anti-chicken AF488 or AF647 (Thermo Fisher A-11039, A-21449) for 20 minutes. After secondary incubation, yeast were washed 2x with PBS + 0.1% BSA and resuspended in a final volume of 1-5 mL of PBS + 0.1% BSA depending on the number of yeast cells stained. When maintaining a high library diversity, all cells showing any binding were sorted. To select for high-affinity clones, the top 1-5% of yeast inside a diagonal gate with increased binding to target peptides proportional to increased scFv expression were sorted into 5 mL of SDCAA media + 1% penicillin-streptomycin and cultured at 30°C at 250 rpm for further selection or diversification processes.

Error-prone PCR was used to affinity mature enriched yeast libraries [68]. Plasmid DNA was extracted from these libraries using the Zymoprep Yeast Plasmid Miniprep II kit (Zymo Research). The antibody region was then PCR-amplified and diversified through EP PCR, consisting of 15-20 cycles with mutagenic nucleotides 8-oxo-dGTP and dPTP (TriLink O-0111, N-2037). The pCTCON2 system primers used were forward, 5’- CGACGATTGAAGGTAGATACCCATACGACGTTCCAGACTACGCTCTGCAG-3’, and reverse, 5’-CGAGCTATTACAAGTCTTCTTCAGAAATAAGCTTTTGTTCTAGAATTCCGGA-3’.

Sequencing primers for pCTCON2 were forward, 5’-GTTCCAGACTACGCTCTGCAGG-3’, and reverse, 5’-GATTTTGTTACATCTACACTGTTG-3’. Second generation (g2.0) libraries were assembled by homologous recombination following co-electroporation of these inserts along with a linearized expression pCTCON2 plasmid containing homologous 5’ and 3’ sequences back into EBY100 *Saccharomyces cerevisiae* yeast as previously described [67]. Sufficiently enriched libraries or g1.0 yeast in the case of the isotype control antibody were plated on SGCAA plates and single colonies were picked and cultured in SGCAA media followed by characterization using the same flow cytometry workflow as previously described for FACS. Clones exhibiting high mutant peptide binding and where possible, low wild-type peptide binding, were chosen for further analysis.

### Isolation of neoepitope-specific antibody fragments by phage display

Three rounds of panning were conducted with either peptide alone or with peptide conjugations to BSA, BSA-biotin, or KLH at Sanyou Biopharmaceuticals. Panning was conducted with a nonimmune, human, Fab-expressing phage library (diversity ≈ 1x10^11^) using both solid-phase and liquid-phase panning methods. In the solid-phase panning method, immunotubes were coated with antigen and blocked with 5% PBSM overnight at 4°C. Phage suspensions, blocked with 5% PBSM, were incubated in the antigen-coated immunotubes.

Binding phages were eluted using trypsin, collected, and used to infect SS320 competent cells spread on selective media plates and incubated overnight at 37°C. Phage titer tests were conducted by diluting the eluted phage solution in a 10-fold gradient and incubating with SS320 cells. Three rounds of panning were performed with decreasing antigen concentration to enrich high-affinity clones. Enrichment was evaluated based on input and output titers.

In the liquid-phase panning method, non-specific phages were depleted using Dynabeads (Thermo Fisher) blocked with 2.5% BSA. Blocked beads were incubated with depleted phages, followed by elution with trypsin. Eluted phages were processed and quantified as described above for each of three rounds of panning with increasing stringency to enrich high-affinity antibody clones.

Positive pool validation was performed by diluting the phage culture supernatant in 5-fold serial dilutions with PBSM (1x PBS, 0.5% bovine serum albumin, 2.5 mM EDTA) and evaluated by ELISA. For the ELISA method, plates were coated with 2 µg/mL of neoepitope peptide, 30 µL/well, overnight at 4°C, then washed with PBST. Plates were blocked with 5% PBSM, 2.5% BSA, or 1% casein at room temperature for 1 hour and washed with PBST. Dilute phage were added (30 µL/well) and incubated at room temperature for 1 hour, then washed with PBST. The secondary antibody, Anti-M13-HRP (phage-specific), was added, incubated at room temperature for 1 hour, and washed with PBST. TMB was added for 5-10 minutes at room temperature. The reaction was stopped with a stop solution, and OD values were read at 450 nm using a microplate reader.

Based on ELISA screening, 644 clones were obtained, of which 162 positive clones were selected for sequencing, resulting in 102 unique clones. From these, 36 clones were selected for full-length construction based on diversity analysis.

### Antibody cloning and expression

Plasmid DNA was extracted from single yeast clones using the Zymoprep Yeast Plasmid Miniprep II kit. Peptide-specific scFv sequences were PCR-amplified (forward, 5’ gcctttctctccacaggcgccatggccCAGGTCCAGCTGGTACAGTC 3’, and reverse 5’ gcggccgcagacaagacccacacctgGGAGAGGACGGTCAGCTGG 3’, lowercase denotes homology with vector, uppercase denotes antibody sequence) and subsequently cloned into a pCMV vector using HiFi assembly (NEBuilder® HiFi DNA Assembly) to reformat these constructs as scFv-Fc antibodies. In the case of the YSD-derived antibodies, the GASDALIE mutations (Gly236Ala/Ser239Asp/Ala330Leu/Ile332Glu) mutations [32–36] were incorporated via site-directed mutagenesis (NEB Q5® Site-Directed Mutagenesis) according to the manufacturer’s protocol. For Fc enhancement of phage-derived antibodies, gene fragments were synthesized (Genewiz) with the GASDALIE mutations as full length human IgG1 kappa antibodies and cloned into a pCMV vector. Unmodified phage-derived antibodies lacking enhancing mutations were produced at Sanyou Bio by PCR-amplifying antibody DNA from phage display vectors and cloning into pcDNA3.4 vectors.

Antibodies were transiently expressed in either Expi293 or ExpiCHO cell expression systems (Thermo Fisher) for 6 days or 8 to 12 days, respectively. Routine culturing occurred in Expi293™ Expression Medium or ExpiCHO™ Expression Medium (Thermo Fisher) Routine culturing and transfections occurred at 37°C, 8% CO_2_, 125 rpm. Certain ExpiCHO transfection protocols required a shift to 32°C 8% CO_2_. Transfection was performed using either Expifectamine^TM^ or PEI as a transfection reagent, with enhancers (Thermo Fisher) added 18-20 hours post-transfection. The antibodies were harvested by first centrifuging at 6,000 g at 4°C for 30-120 minutes followed by filtering through 0.45 µm surfactant-free cellulose acetate filters.

Antibodies were then purified using 1- to 2-mL of protein A or G resin (Genscript L00210 and L00209) with gravity columns (Bio-Rad 7321010). An additional wash step was conducted while the antibody was bound to the column using 6 column volumes of 100 mM sodium carbonate, pH 10.5, to improve antibody quality [69]. Elution was performed using 10 mL of 100 mM glycine, pH 3.0 followed by 2 mL of 100 mM glycine, pH 2.7 directly into 1 mL of 1M TRIS-HCl, pH 8 neutralization buffer (Teknova T1080). Subsequently, antibodies were filtered through 0.22 µm PVDF filters, and their concentrations were determined at A280 using each antibody’s extinction coefficient calculated from their sequences (Expasy ProtParam tool).

### Peptide binding assays

To evaluate binding of reformatted antibodies, biotinylated peptides were conjugated to SA-coated beads as described above. Candidate mAbs diluted in PBS + 0.1% BSA were incubated with these peptide-coated beads for at least 1 hour at 4°C with shaking at 900 rpm, followed by 2x washes with PBS + 0.1% BSA and by incubation with a mouse anti-human IgG Fc-PE secondary antibody (SouthernBiotech 9040-09) at 1 µg/mL for 20-30 minutes at 4°C with shaking at 900 rpm. Finally, the beads were washed 2x with PBS + 0.1% BSA before quantification of antibody binding using an Agilent NovoCyte Advanteon flow cytometer. Data was analyzed as PE median fluorescence intensity (MFI) after gating out doublets. Binding of the phage-derived antibodies to peptide conjugates was quantified by EC_50_ values using ELISA at Sanyou Biopharmaceuticals.

### Tumor and effector cell culture

B16F10 (ATCC CRL-6475), EMT6 (ATCC CRL-2755), A375 (ATCC CRL-1619), and PANC1 (ATCC CRL-1469) cells were cultured in DMEM supplemented with 10% FBS, 100 U/mL penicillin, and 100 µg/mL streptomycin at 37°C, 5% CO_2_. B16F10 cells used in the *in vivo* models were cultured in RPMI supplemented with 7.5% FBS. Jurkat-Lucia NFAT-CD16 reporter cells (InvivoGen jktl-nfat-cd16) were cultured in DMEM supplemented with 10% FBS, 100 U/mL penicillin, 100 µg/mL streptomycin, 100 µg/mL Normocin, 10 µg/mL of Blasticidin, and 100 µg/mL of Zeocin. THP-1 cells (ATCC TIB-202) were cultured in RPMI supplemented with 10% FBS, 0.05 mM 2-mercaptoethanol, 100 U/mL penicillin, and 100 µg/mL streptomycin. CT26 cells (ATCC CRL-2638) were cultured in RPMI supplemented with 10% FBS, 100 U/mL penicillin, and 100 µg/mL streptomycin.

### Cell binding assays

For cell binding assays, B16F10 cells were lifted from tissue culture treated flasks using 2.5 mM EDTA to preserve cell surface proteins and washed with PBS. Cells were incubated with neoepitope-targeting mAbs and subsequently detected with a mouse anti-human IgG Fc-PE secondary antibody (SouthernBiotech 9040-09). Detection of the rabbit pAb cocktail was conducted with a goat anti-rabbit IgG (H+L) AF647 secondary antibody (Abcam ab150079). Flow cytometry was used for quantifying binding using an Agilent NovoCyte Advanteon flow cytometer.

Antibody binding to cells was also visualized via fluorescence microscopy using a Nikon Spinning Disk Confocal Yokogawa CSU. Single cell suspensions (30 µL containing 20,000 cells) were added per well of µ-Slide VI 0.4 culture slides (Ibidi 80606). Cells were allowed to adhere for 30 minutes at 37°C, 5% CO_2_ under humid conditions before adding fresh media for overnight growth. The slides were then washed 3x with 100 µL PBS and fixed with 100 µL 4% paraformaldehyde for 15 min at RT. Slides were then washed 3x with 100 µL PBS and incubated 100 µL with Image-iT™ FX Signal Enhancer (Thermo Fisher) for 30 min at RT. Slides were then washed 3x with 100 µL PBS and blocked with PBS + 1% BSA for 1 hour at RT. Slides were stained with 100 µL of the primary antibody diluted in PBS + 1% BSA overnight at 4°C. The following day, slides were allowed to warm to RT, washed 3x with PBS + 0.1% BSA, and incubated with a goat anti-human IgG (H+L) AF647 secondary antibody (Thermo Fisher A- 21445) for 1 hour at RT in the dark. Slides were washed 6x with 100 µL PBS and then incubated with DAPI nuclear stain (Thermo Fisher D3571) diluted 1:1000 in sterile water for 10 min at RT in the dark. Incubation steps took place in a moist box. Finally, slides were washed 3x with 100 µL PBS before imaging. Quantification of staining intensity was measured by randomly selecting three or more microscopic fields of view, subtracting the background, which was defined by measuring the intensity of a blank area of the image with no cells or debris, and calculating the ratio of antibody staining intensity to DAPI intensity to account for cell number in each field.

Antibody binding to cells was also visualized via fluorescence microscopy in the context of tissues using a Thermo Fisher EVOS M5000. B16F10 tissues were harvested from B16F10- bearing C57BL/6 mice while wild-type tissues were harvested from naïve C57BL/6 mice. These tissues were formalin fixed and frozen before added to microscope slides for staining. The slides were thawed for 3 minutes at RT and fixed with 3% paraformaldehyde for 15 min at RT. The slides were then washed for 3 minutes 2x with PBS followed by blocking with 100 µL of 5% normal goat serum block diluted in PBS + 1.0% BSA + 0.1% Triton X-100 for 1 hour at RT. Slides were then washed for 5 minutes 3x with PBS at RT. 100 µL of neoepitope-targeting antibodies diluted in PBS + 1.0% BSA were then incubated with the slides overnight at 4°C. Slides were then washed for 5 minutes 7x with PBS 1.0% BSA at RT followed by detection with a goat anti-human IgG (H+L) AF488 secondary antibody (Thermo Fisher A-11013) diluted in PBS + 1.0% BSA for 1 hour at RT. Slides were then washed for 5 minutes 3x with PBS 1.0% BSA at RT. Finally, slides were incubated with DAPI nuclear stain (Thermo Fisher D3571) diluted 1:1000 in sterile water for 10 minutes at RT in the dark, followed by three washes for 5 minutes each with PBS + 1% BSA at RT before mounting with DAKO fluorescent mounting medium (Agilent S3023) and imaging. Incubation steps took place in a moist box. Quantification of staining intensity was measured by randomly selecting three or more microscopic fields of view, subtracting the background, which was defined by measuring the intensity of a blank area of the image with no cells or debris, and calculating the ratio of antibody staining intensity to DAPI intensity to account for cell number in each field.

Immunocytochemistry staining of B16F10 cells with phage-derived antibodies was conducted at Sanyou Biopharmaceuticals. Briefly, B16F10 cells were seeded in 12-well plates 48 hours prior to staining and fixed with 4% paraformaldehyde upon reaching 80% confluence. Permeabilization was performed using 0.5% Triton X-100 for 20 minutes. To block endogenous enzyme activity, cells were incubated with 3% hydrogen peroxide for 15 minutes. Phage-derived antibody, at a concentration of 5 µg/mL, was incubated with the cells at 37°C for 60 minutes or at 4°C overnight. Secondary detection was carried out using a Goat Anti-Human IgG Fc-HRP (Abcam ab97225) at a 1:1000 dilution, with incubation at 37°C for 30 minutes. This was followed by 3,3’-diaminobenzidine chromogen staining at 37°C for 30 minutes. Imaging was performed using a Zeiss microscope.

### Effector function assays

ADCC was assessed using a proxy signaling assay that employs Jurkat-Lucia NFAT- CD16 reporter cells (InvivoGen). These reporter cells emit a luminescent signal upon engagement of FcγRIIIa (CD16) receptors, providing a quantitative measure of FcγRIIIa ligation and activation. In 96-well tissue culture treated plates, 20,000 live B16F10 cells and 80,000 live Jurkat reporter cells were added to each well immediately one after another in culture media (RPMI supplemented with 10% FBS, 1 mM sodium pyruvate, 1x non-essential amino acids, and 100 U/mL penicillin, and 100 µg/mL streptomycin) followed by antibody diluted in culture media for a total of 200 µL. Plates were then incubated for 24 or 48 hours at 37°C, 5% CO_2_. A cell stimulation cocktail (Thermo Fisher 00-4970-93) was used to induce cell activation as a positive control. Plates were then centrifuged for 5 min at 500 g and 80 µL of the supernatant was added to a different opaque 96-well plates containing 20 µL of QUANTI-Luc™ Luciferase Detection Reagent (InvivoGen). With the lid off, data was immediately acquired on a Molecular Devices SpectraMax Paradigm plate reader to detect luminescence in each well. The relative light units (RLU) average of three measurements spaced 2.5 minutes apart was used for assessing differences in ADCC potential.

ADCP was evaluated by monitoring the phagocytosis of B16F10 cells by THP-1 monocytes. For this assay, B16F10 cells were stained with CellTracker™ Deep Red (Thermo Fisher), and THP-1 monocytes were labeled with CellTrace™ CFSE (Thermo Fisher) according to the manufacture’s protocols. A total of 100,000 stained B16F10 cells and 100,000 stained THP-1 cells were added to non-tissue culture treated 96-well plates and incubated for 4 hours with mild shaking (400 rpm) at 37°C, 5% CO_2_. Antibodies were used at a concentration of 20 µg/mL. Flow cytometry was used to measure the phagocytosis score, which was calculated as the proportion of labeled cells that were double positive among the total THP-1 cells, multiplied by the MFI of the THP-1 cells. Additionally, fluorescence microscopy (Microscope: Nikon Spinning Disk Confocal Yokogawa CSU) was employed to visually confirm and analyze this phagocytic activity. The isotype control scFv-Fc antibody served as a negative control in these experiments.

### *In vivo* mouse models

Female C57BL/6 mice, 8 - 12 weeks old, were obtained from Charles River Laboratories (Sanyou Bio experiments) or Jackson Laboratory (Dartmouth experiments) and housed within specific pathogen-free rooms. Up to five mice were co-housed per isolator caging unit in a temperature-regulated environment with standard light and dark cycles. Studies were performed with approval from Dartmouth’s Institutional Animal Care and Use Committee.

Antibodies underwent endotoxin removal (Thermo Fisher Pierce™ High Capacity Endotoxin Removal spin columns) followed by subsequent filtering using 0.22 µm PVDF filters prior to animal administration. Endotoxin levels of all antibodies were measured to be below 2 EU/mL (GenScript ToxinSensor™ Chromogenic LAL Endotoxin Assay Kit).

The ICB antibodies used were rat anti-mouse PD-1 (Bio X Cell BE0146, Clone RMP1- 14) and mouse anti-mouse CTLA-4 (Bio X Cell BE0164, Clone 9D9). The IgG control used in *in vivo* experiments was an HIV-specific antibody, VRC01.

Depending on the experiment, 5x10^5^ (Sanyou Bio experiments) or 3x10^5^ (Dartmouth experiments) B16F10 cells lifted from tissue culture treated flasks with Trypsin (Corning 25-052- CI) were implanted intradermally on the shaved right flank of each mouse. Depending on the experiment, three (Sanyou Bio experiments) or five (Dartmouth experiments) days post tumor implantation, mice with confirmed tumors were grouped based on the average tumor size per group. Mice without tumors on the day of grouping were excluded from study. Anti-B16F10 antibody treatments were administered intradermally at the base of the tumor, while ICB antibodies were administered intraperitoneally. Antibodies were administered in a volume of 100 µL, containing either 200 or 800 µg of B16F10-targeting antibodies. Each ICB antibody was administered at a dose of 200 µg. Treatment schema can be found in the figures of each *in vivo* experiment. For the Sanyou Bio experiments, tumor volumes were measured using mechanical calipers and the following formula: L x W^2^ / 2. For the Dartmouth experiments, mechanical calipers were used to take measurements in an "X" pattern, measuring the width and length of the tumor, followed by a measurement of the tumor height to calculate the tumor volume (L x W x H). Measurements were taken before treatment if a measurement day overlapped with a treatment day. Mouse weight was monitored throughout the experiments. Mice were sacrificed in accordance with the local Institutional Animal Care and Use Committee Guidance.

### Software tools and analytical methods for data processing

Statistical analyses were conducted in GraphPad Prism version 10.1.2 for Windows. Details about which tests were run can be found in the figure legends. Microsoft Excel was utilized for data arrangement, experiment planning, and calculating treatment synergy with the Bliss Independence Model [29]. Flow cytometric analyses were conducted in Flowlogic version 8.7 for Windows. Fluorescent microscopy images were analyzed in ImageJ version 1.53 for Windows.

### Handling conflicts of interest

All authors and collaborators fully disclosed any potential conflicts of interest, including financial, personal, and professional relationships that could potentially influence the research outcomes. Data collection, analysis, and interpretation were conducted objectively, with all authors adhering to ethical research practices.

---

**Supplemental Figure 1.**
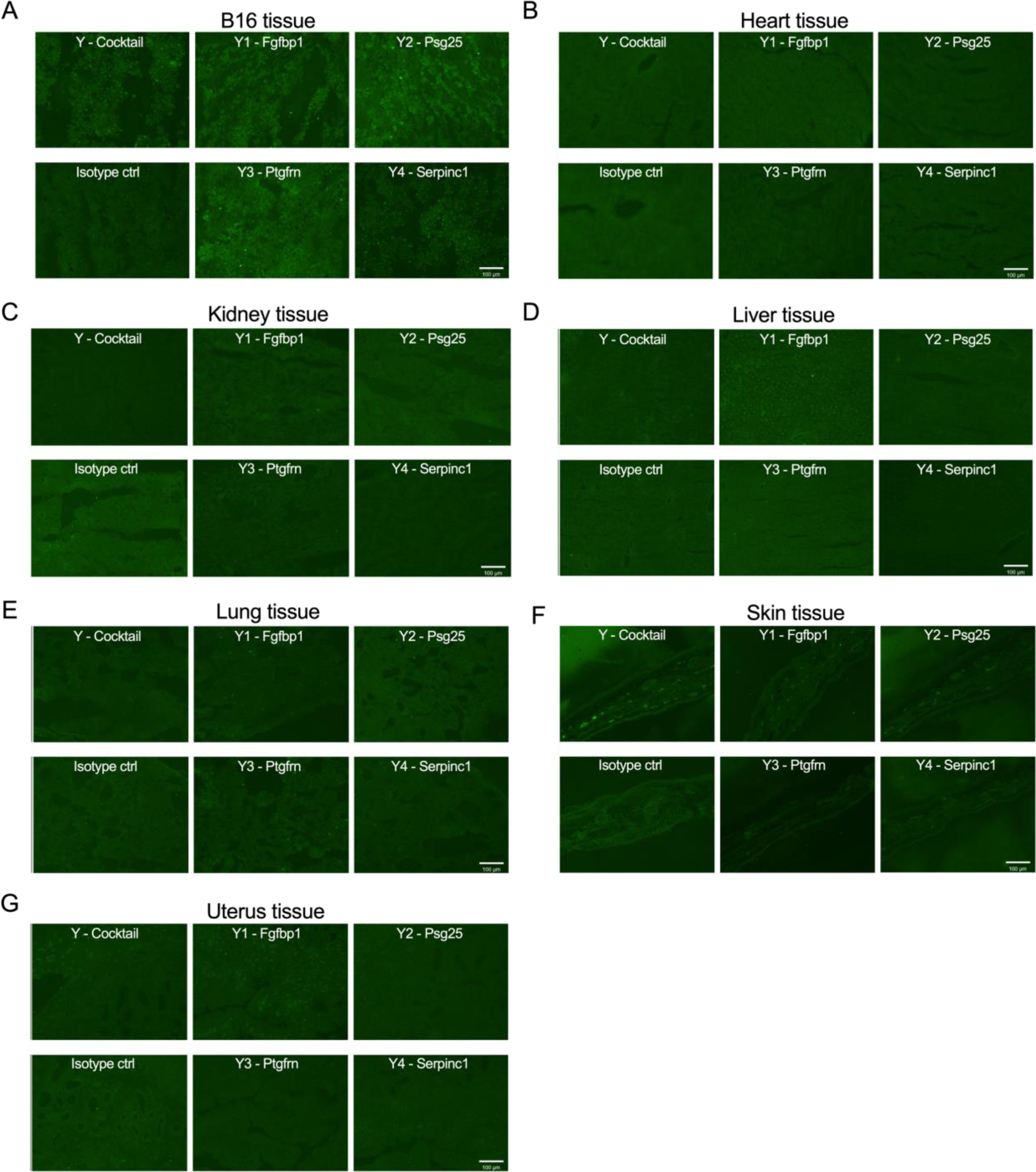
On- and off-target binding of the yeast-derived antibodies to a tissue panel. **A-G.** Fluorescence microscopy staining of malignant B16F10 tissue (**A**) and a panel of wild-type C57BL/6 tissues form non-tumor-bearing mice including heart (**B**), kidney (**C**), liver (**D**), lung (**E**), skin (**F**), and uterus (**G**).

**Supplemental Figure 2.**
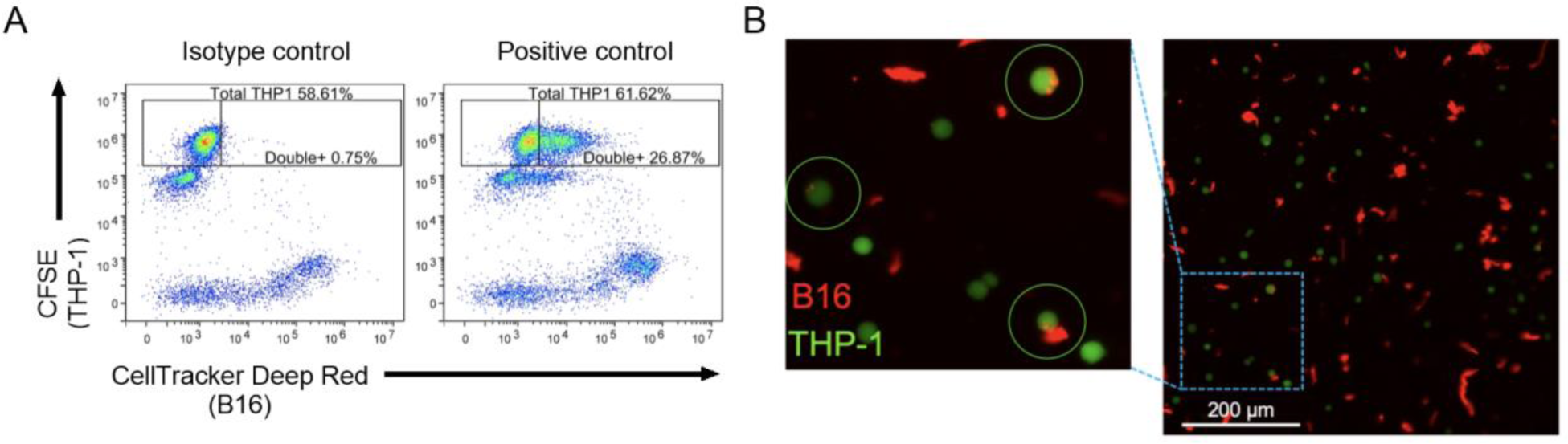
Antibody-dependent phagocytosis assay with THP-1 monocytes and B16F10 cells. **A.** Exemplary negative (left) and positive (right) phagocytosis assay flow biplots demonstrating phagocytosis/trogocytosis of B16F10 cells/membrane by THP-1 monocytes. **B.** Fluorescent microscopy staining highlighting trogocytosis activity of THP-1 monocytes against B16F10 cells in the presence of B16F10-specific antibody. CFSE, carboxyfluorescein diacetate succinimidyl ester.

**Supplemental Figure 3.**
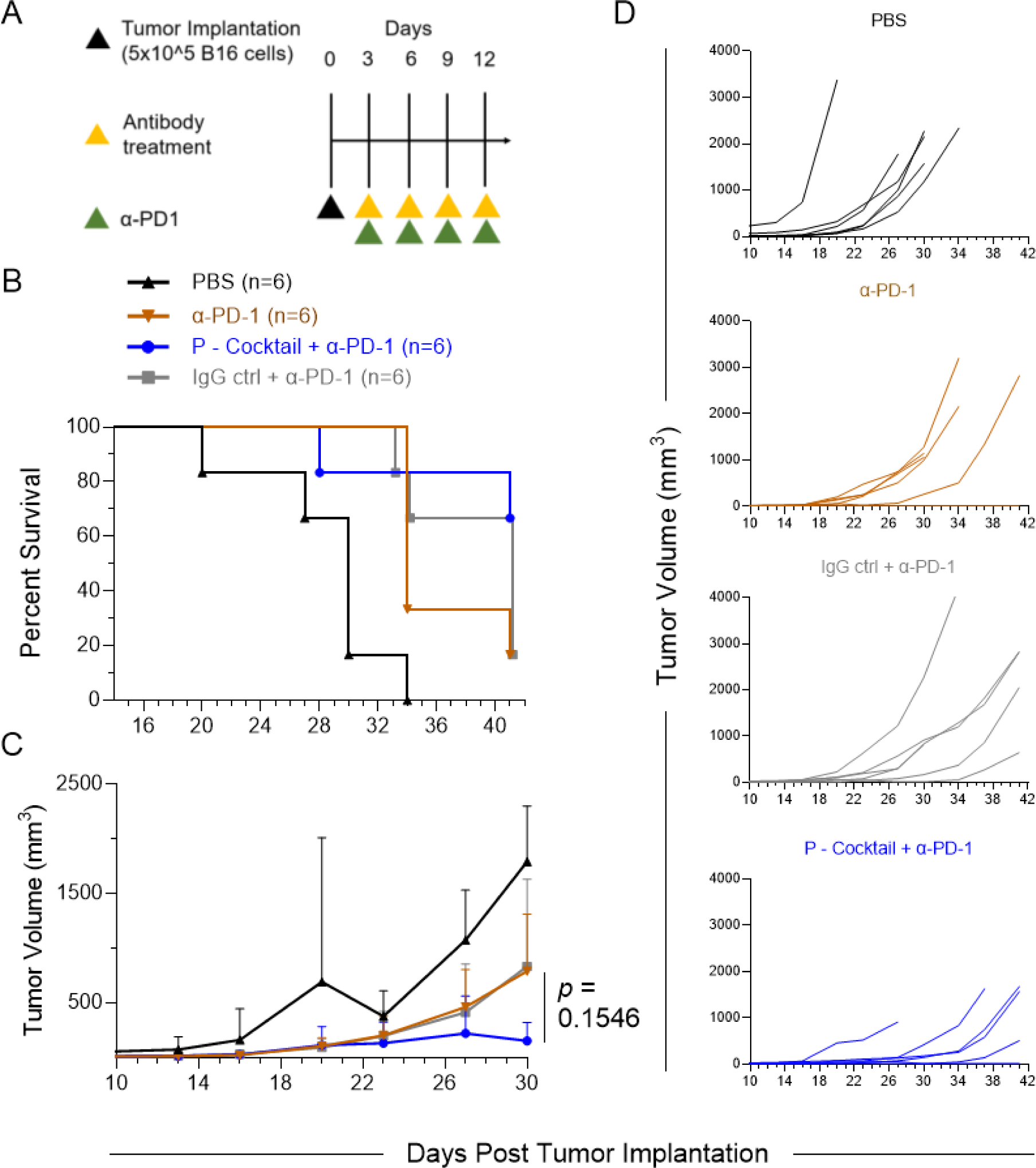
Treatment of B16F10-bearing C57BL/6J mice with neoepitope- targeted phage-derived antibody cocktail. **A.** Treatment regimen. **B.** Kaplan-Meier survival curves. A log-rank Mantel-Cox test was used to calculate a difference in the survival curves (*p* ≥ 0.05). **C.** Tumor growth curves are depicted over time following confirmed tumor implantation. A two-way ANOVA (Time: *F*_1.264, 18.75_ = 16.99, *p* = 0.0003; Treatment: *F*_2, 15_ = 1.104, *p* = 0.3570; Time x Treatment: *F*_12, 89_ = 1.929, *p* = 0.0411) with a Greenhouse-Geisser correction (ε = 0.2106) followed by Dunnett’s post hoc test was used to compare P - Cocktail + α -PD-1 against the IgG control + α-PD-1 group. Error bars represent SD of the mean. **D.** Tumor growth curves for individual mice.

**Supplemental Figure 4.**
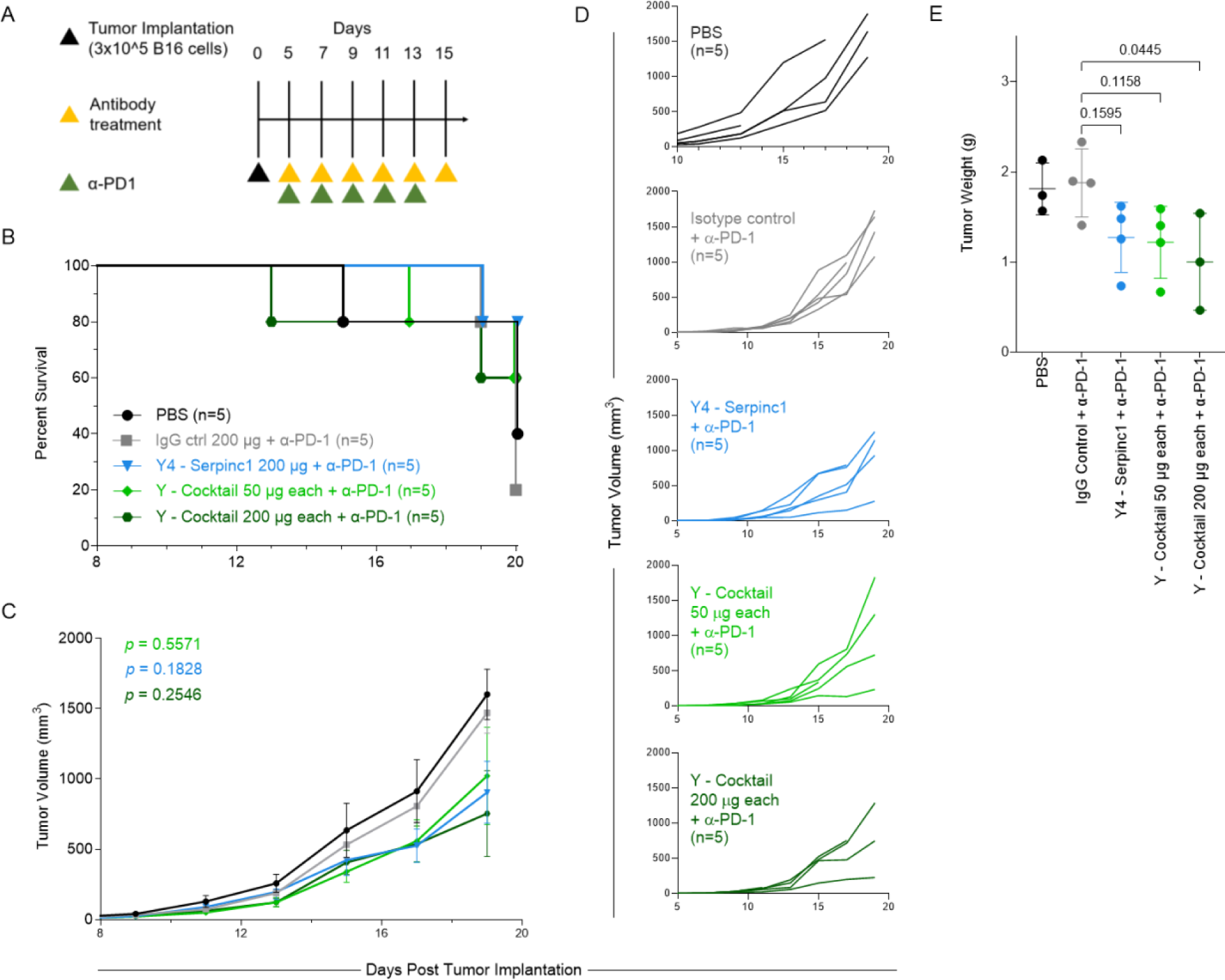
Treatment of B16F10-bearing C57BL/6J mice with neoepitope- targeted yeast-derived antibody cocktail. **A.** Treatment regimen. **B.** Kaplan-Meier survival curves. A log-rank Mantel-Cox test was used to calculate a difference in the survival curves (*p* ≥ 0.05). **C**. Tumor growth curves are depicted over time following confirmed tumor implantation. A two-way ANOVA (Time: *F*_1.246, 18.54_ = 81.45, *p* < 0.0001; Treatment: *F*_3, 16_ = 1.693, *p* = 0.2087; Time x Treatment: *F*_24, 119_ = 1.644, *p* = 0.0428) with a Greenhouse-Geisser correction (ε = 0.1558) followed by Dunnett’s post hoc test was used to compare treatment groups against the IgG control + α-PD-1 group. Error bars represent SD of the mean. **D.** Tumor growth curves for individual mice. **E.** Weights of excised tumors from all surviving mice on day 20 at the termination of the experiment. A one-way ANOVA followed by Dunnett’s post hoc test was used to compare treatment groups against the IgG control + α-PD-1 group.

**Supplemental Figure 5.**
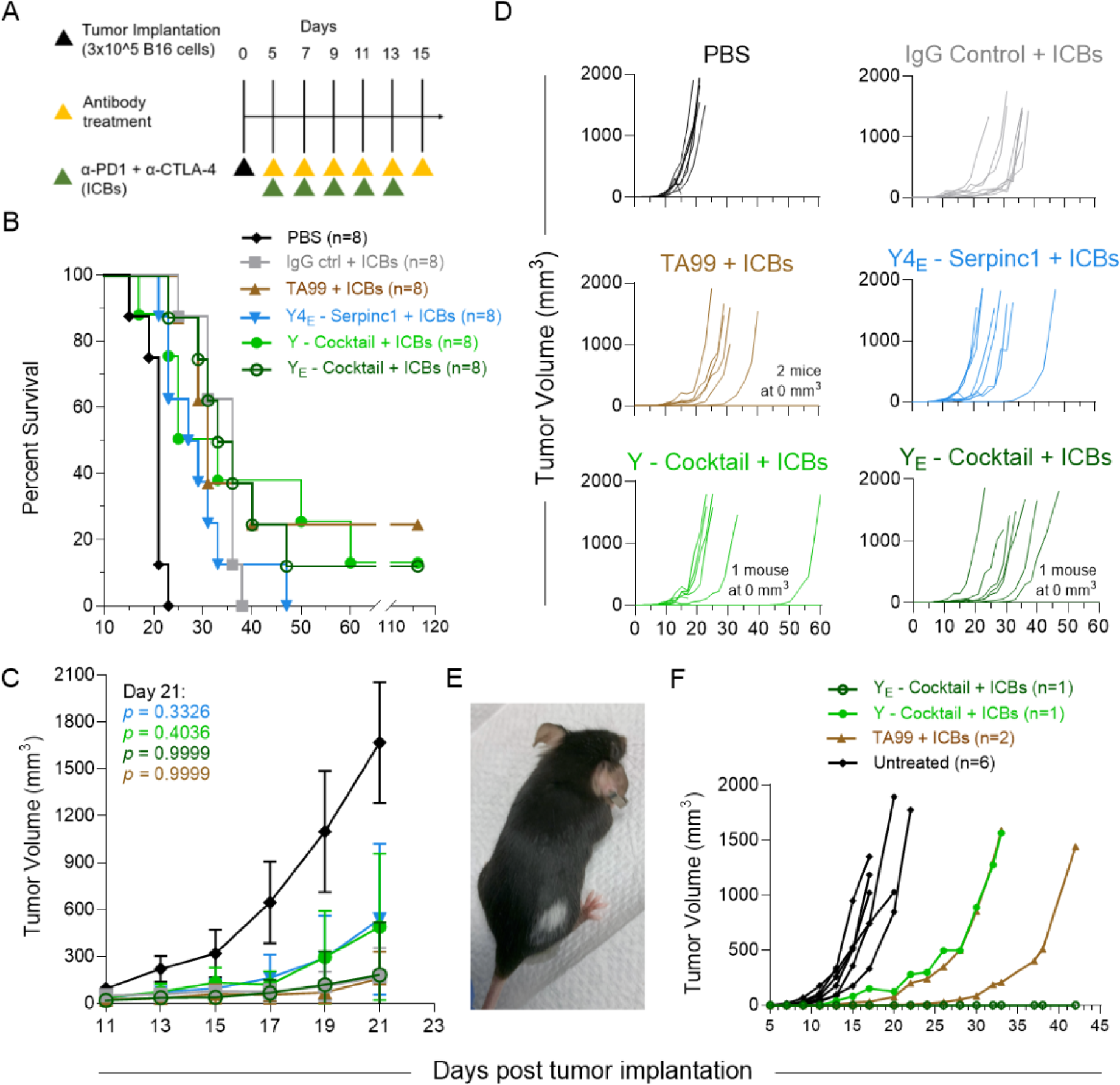
Treatment and re-challenge of B16F10-bearing C57BL/6J mice with neoepitope-targeted yeast-derived antibody cocktail. **A.** Treatment regimen in which individual antibodies, antibody cocktails, or isotype control and immune checkpoint blockade (ICBs) or buffer control (PBS) were administered to tumor-bearing mice. **B.** Kaplan-Meier survival curves. A log-rank Mantel-Cox test was used to calculate a difference in the survival curves (*p* ≥ 0.05). TA99 served as a positive control antibody known to be effective in B16- bearing mice. **C.** Tumor growth curves over time following confirmed tumor implantation. A two- way ANOVA (Time: *F*_1.087, 37.82_ = 20.46, *p* < 0.0001; Treatment: *F*_4, 35_ = 1.672, *p* = 0.1786; Time x Treatment: *F*_36, 313_ = 1.698, *p* = 0.0096) with a Greenhouse-Geisser correction (ε = 0.1208) followed by Dunnett’s post hoc test was used to compare treatment groups against the IgG control + ICBs group (*p* ≥ 0.05). Error bars represent SD of the mean. **D.** Tumor growth curves for individual mice. **E.** Image of the surviving mouse from the Y - Cocktail + ICBs group presenting with vitiligo. **F.** Tumor growth curves from the re-challenge of individual surviving mice with 3x10^5^ B16F10 cells on the same right flank 117 days post initial tumor implantation and untreated naïve control mice.

**Supplemental Figure 6.**
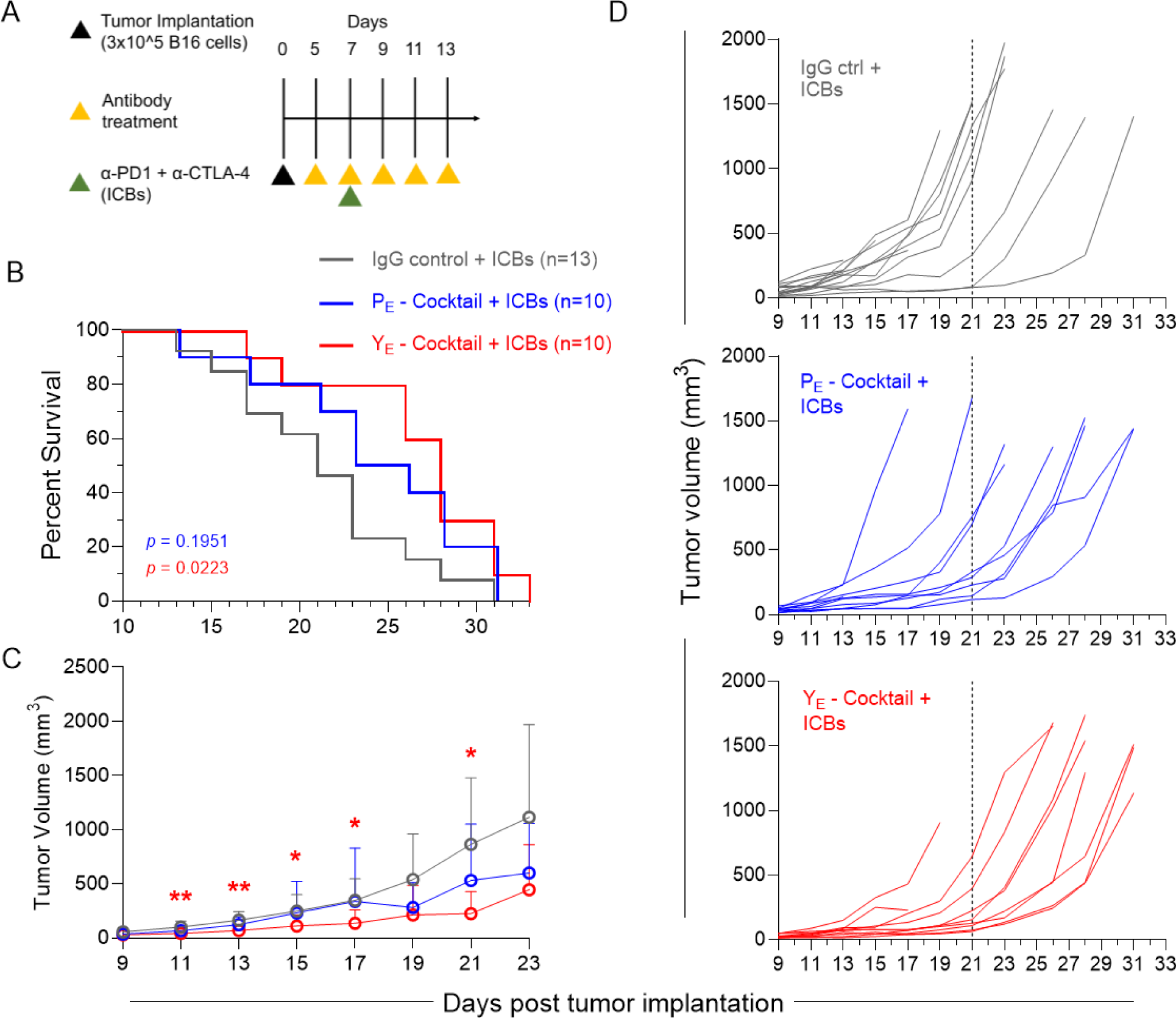
Repeat experiment of neoepitope-targeted yeast- and phage-derived antibody cocktails in B16F10-bearing C57BL/6J mice. **A.** Treatment regimen in which antibody cocktails or isotype control (IgG ctrl) and immune checkpoint blockade (ICBs) were administered to tumor-bearing mice. **B.** Kaplan-Meier survival curves. A log-rank Mantel-Cox test was used to calculate a difference in the survival curves for phage and yeast antibody cocktails as compared to isotype control. **C.** Tumor growth curves are depicted over time following confirmed tumor implantation. A two-way ANOVA (Time: *F3.691*, _82.33_ = 47.63, *p* < 0.0001; Treatment: *F*_2, 30_ = 3.960, *p* = 0.0298; Time x Treatment: *F*_26, 290_ = 1.891, *p* = 0.0066) with a Greenhouse-Geisser correction (ε = 0.2839) followed by Dunnett’s post hoc test was used to compare treatment groups against the IgG control + ICBs group. Error bars represent SD of the mean, * *p* < 0.05, ** *p* < 0.01. **D.** Tumor growth curves for individual mice. The vertical line indicates the median survival of mice in the IgG control + ICBs group.

## REFERENCES

1. Shukla GS, Sun YJ, Pero SC, et al. A cocktail of polyclonal affinity enriched antibodies against melanoma mutations increases binding and inhibits tumor growth. J Immunol Methods. 2020;478:112720.

2. Varghese B, Widman A, Do J, et al. Generation of CD8+ T cell-mediated immunity against idiotype-negative lymphoma escapees. Blood. 2009;114(20):4477–85.

3. Gambardella V, Tarazona N, Cejalvo JM, et al. Personalized Medicine: Recent Progress in Cancer Therapy. Cancers (Basel). 2020;12(4).

4. Cai WQ, Zeng LS, Wang LF, et al. The Latest Battles Between EGFR Monoclonal Antibodies and Resistant Tumor Cells. Front Oncol. 2020;10:1249.

5. James D Mellow MPB, Helen R Irving, John R Zalcberg, Alexander Dobrovic. A critical review of the role of Fc gamma receptor polymorphisms in the response to monoclonal antibodies in cancer. Journal of Hematology & Oncology. 2013;6.

6. Jiang T, Shi T, Zhang H, et al. Tumor neoantigens: from basic research to clinical applications. Journal of Hematology & Oncology. 2019;12(1).

7. Blass E, Ott PA. Advances in the development of personalized neoantigen-based therapeutic cancer vaccines. Nat Rev Clin Oncol. 2021;18(4):215–29.

8. Shukla GS, Sun YJ, Pero SC, et al. Immunization with tumor neoantigens displayed on T7 phage nanoparticles elicits plasma antibody and vaccine-draining lymph node B cell responses. J Immunol Methods. 2018;460:51–62.

9. Richard G, Princiotta MF, Bridon D, et al. Neoantigen-based personalized cancer vaccines: the emergence of precision cancer immunotherapy. Expert Rev Vaccines. 2022;21(2):173–84.

10. Wirth TC, Kuhnel F. Neoantigen Targeting-Dawn of a New Era in Cancer Immunotherapy? Front Immunol. 2017;8:1848.

11. Linnemann C, van Buuren MM, Bies L, et al. High-throughput epitope discovery reveals frequent recognition of neo-antigens by CD4+ T cells in human melanoma. Nat Med. 2015;21(1):81–5.

12. Xie N, Shen G, Gao W, et al. Neoantigens: promising targets for cancer therapy. Signal Transduct Target Ther. 2023;8(1):9.

13. Zhang S, Helling, F., Lloyd, K O., Livingston, P O. Increased tumor cell reactivity and complement-dependent cytotoxicity with mixtures of monoclonal antibodies against different gangliosides. Cancer Immunology Immunotherapy. 1995;40(2).

14. Glassy MC, McKnight ME, Kotlan B, et al. Cocktails of human anti-cancer antibodies show a synergistic effect in nude mouse tumor xenografts. Human Antibodies. 2008;16(3-4):87–98.

15. Glassy MC, McKnight ME. Requirements for human antibody cocktails for oncology. Expert Opin Biol Ther. 2005;5(10).

16. Tran E, Turcotte S, Gros A, et al. Cancer immunotherapy based on mutation-specific CD4+ T cells in a patient with epithelial cancer. Science. 2014;344(6184):641-5.

17. Geukes Foppen MH, Donia M, Svane IM, et al. Tumor-infiltrating lymphocytes for the treatment of metastatic cancer. Mol Oncol. 2015;9(10):1918–35.

18. Yadav M, Jhunjhunwala S, Phung QT, et al. Predicting immunogenic tumour mutations by combining mass spectrometry and exome sequencing. Nature. 2014;515(7528):572-6.

19. Weber JS, Carlino MS, Khattak A, et al. Individualised neoantigen therapy mRNA-4157 (V940) plus pembrolizumab versus pembrolizumab monotherapy in resected melanoma (KEYNOTE-942): a randomised, phase 2b study. Lancet. 2024;403(10427):632–44.

20. Rojas LA, Sethna Z, Soares KC, et al. Personalized RNA neoantigen vaccines stimulate T cells in pancreatic cancer. Nature. 2023;618(7963):144-50.

21. Peng M, Mo Y, Wang Y, et al. Neoantigen vaccine: an emerging tumor immunotherapy. Mol Cancer. 2019;18(1):128.

22. Miller RA. Treatment of B-cell lymphoma with monoclonal anti-idiotype antibody. The New England Journal of Medicine. 1982;306:517–22.

23. Maloney DG, Brown S, Czerwinski DK, et al. Monoclonal anti-idiotype antibody therapy of B-cell lymphoma: the addition of a short course of chemotherapy does not interfere with the antitumor effect nor prevent the emergence of idiotype-negative variant cells. Blood. 1992;80(6):1502–10.

24. Shukla GS, Pero SC, Sun Y, et al. Multiple antibodies targeting tumor-specific mutations redirect immune cells to inhibit tumor growth and increase survival in experimental animal models. Clin Transl Oncol. 2020;22(7):1094–104.

25. Feldhaus MJ, Siegel RW, Opresko LK, et al. Flow-cytometric isolation of human antibodies from a nonimmune Saccharomyces cerevisiae surface display library. Nat Biotechnol. 2003;21(2):163–70.

26. Castle JC, Kreiter S, Diekmann J, et al. Exploiting the mutanome for tumor vaccination. Cancer Res. 2012;72(5):1081–91.

27. Hong Y, Guo H, Wei M, et al. Cell-based reporter assays for measurements of antibody-mediated cellular cytotoxicity and phagocytosis against SARS-CoV-2 spike protein. J Virol Methods. 2022;307:114564.

28. Talathi SP, Shaikh NN, Pandey SS, et al. FcgammaRIIIa receptor polymorphism influences NK cell mediated ADCC activity against HIV. BMC Infect Dis. 2019;19(1):1053.

29. Liu Q, Yin X, Languino LR, et al. Evaluation of drug combination effect using a Bliss independence dose-response surface model. Stat Biopharm Res. 2018;10(2):112–22.

30. Phuangbubpha P, Thara S, Sriboonaied P, et al. Optimizing THP-1 Macrophage Culture for an Immune-Responsive Human Intestinal Model. Cells. 2023;12(10).

31. Nakamura M, Ono D, Sugita S. Mechanophenotyping of B16 Melanoma Cell Variants for the Assessment of the Efficacy of (-)-Epigallocatechin Gallate Treatment Using a Tapered Microfluidic Device. Micromachines (Basel). 2019;10(3).

32. Smith P, DiLillo DJ, Bournazos S, et al. Mouse model recapitulating human Fcgamma receptor structural and functional diversity. Proc Natl Acad Sci U S A. 2012;109(16):6181–6.

33. Bournazos S, Klein F, Pietzsch J, et al. Broadly neutralizing anti-HIV-1 antibodies require Fc effector functions for in vivo activity. Cell. 2014;158(6):1243–53.

34. Ahmed AA, Keremane SR, Vielmetter J, et al. Structural characterization of GASDALIE Fc bound to the activating Fc receptor FcgammaRIIIa. J Struct Biol. 2016;194(1):78–89.

35. Lazar G. Engineered antibody Fc variants with enhanced effector function. PNAS. 2006;103(11):4005–10.

36. Shields RL, Namenuk AK, Hong K, et al. High resolution mapping of the binding site on human IgG1 for Fc gamma RI, Fc gamma RII, Fc gamma RIII, and FcRn and design of IgG1 variants with improved binding to the Fc gamma R. J Biol Chem. 2001;276(9):6591-604.

37. Byrne KT, Cote AL, Zhang P, et al. Autoimmune melanocyte destruction is required for robust CD8+ memory T cell responses to mouse melanoma. J Clin Invest. 2011;121(5):1797–809.

38. Chen D, Xu Z, Cui J, et al. A mouse model of vitiligo based on endogenous auto-reactive CD8 + T cell targeting skin melanocyte. Cell Regen. 2022;11(1):31.

39. Byrne KT, Zhang P, Steinberg SM, et al. Autoimmune vitiligo does not require the ongoing priming of naive CD8 T cells for disease progression or associated protection against melanoma. J Immunol. 2014;192(4):1433–9.

40. Loo K, Smithy JW, Postow MA, et al. Factors Determining Long-Term Antitumor Responses to Immune Checkpoint Blockade Therapy in Melanoma. Front Immunol. 2021;12:810388.

41. Beattie J, Rizvi H, Fuentes P, et al. Success and failure of additional immune modulators in steroid-refractory/resistant pneumonitis related to immune checkpoint blockade. J Immunother Cancer. 2021;9(2).

42. Dobosz P, Stepien M, Golke A, et al. Challenges of the Immunotherapy: Perspectives and Limitations of the Immune Checkpoint Inhibitor Treatment. Int J Mol Sci. 2022;23(5).

43. Miller ML, Finn OJ. Flow cytometry-based assessment of direct-targeting anti-cancer antibody immune effector functions. Methods Enzymol. 2020;632:431–56.

44. Van Wagoner CM, Rivera-Escalera F, Jaimes-Delgadillo NC, et al. Antibody-mediated phagocytosis in cancer immunotherapy. Immunol Rev. 2023;319(1):128–41.

45. Cao X, Chen J, Li B, et al. Promoting antibody-dependent cellular phagocytosis for effective macrophage-based cancer immunotherapy. Sci Adv. 2022;8(11):eabl9171.

46. Ochoa MC, Minute L, Rodriguez I, et al. Antibody-dependent cell cytotoxicity: immunotherapy strategies enhancing effector NK cells. Immunol Cell Biol. 2017;95(4):347–55.

47. Pinto S, Pahl J, Schottelius A, et al. Reimagining antibody-dependent cellular cytotoxicity in cancer: the potential of natural killer cell engagers. Trends Immunol. 2022;43(11):932–46.

48. Wang W, Erbe AK, Hank JA, et al. NK Cell-Mediated Antibody-Dependent Cellular Cytotoxicity in Cancer Immunotherapy. Front Immunol. 2015;6:368.

49. Zahavi D, AlDeghaither D, O’Connell A, et al. Enhancing antibody-dependent cell-mediated cytotoxicity: a strategy for improving antibody-based immunotherapy. Antib Ther. 2018;1(1):7–12.

50. Lara S, Heilig J, Virtanen A, et al. Exploring complement-dependent cytotoxicity by rituximab isotypes in 2D and 3D-cultured B-cell lymphoma. BMC Cancer. 2022;22(1):678.

51. Kolev M, Das M, Gerber M, et al. Inside-Out of Complement in Cancer. Front Immunol. 2022;13:931273.

52. Macor P, Capolla S, Tedesco F. Complement as a Biological Tool to Control Tumor Growth. Front Immunol. 2018;9:2203.

53. Baniel CC, Sumiec EG, Hank JA, et al. Intratumoral injection reduces toxicity and antibody-mediated neutralization of immunocytokine in a mouse melanoma model. J Immunother Cancer. 2020;8(2).

54. Marabelle A, Tselikas L, de Baere T, et al. Intratumoral immunotherapy: using the tumor as the remedy. Ann Oncol. 2017;28(suppl_12):xii33-xii43.

55. Blanco E, Chocarro L, Fernandez-Rubio L, et al. Leading Edge: Intratumor Delivery of Monoclonal Antibodies for the Treatment of Solid Tumors. Int J Mol Sci. 2023;24(3).

56. Li JQ, Zhang ZR, Zhang HQ, et al. Intranasal delivery of replicating mRNA encoding neutralizing antibody against SARS-CoV-2 infection in mice. Signal Transduct Target Ther. 2021;6(1):369.

57. Deal CE, Carfi A, Plante OJ. Advancements in mRNA Encoded Antibodies for Passive Immunotherapy. Vaccines (Basel). 2021;9(2).

58. Rybakova Y, Kowalski PS, Huang Y, et al. mRNA Delivery for Therapeutic Anti-HER2 Antibody Expression In Vivo. Mol Ther. 2019;27(8):1415–23.

59. Martinez-Navio JM, Fuchs SP, Mendes DE, et al. Long-Term Delivery of an Anti-SIV Monoclonal Antibody With AAV. Front Immunol. 2020;11:449.

60. Piperno GM, Lopez-Requena A, Predonzani A, et al. Recombinant AAV-mediated in vivo long-term expression and antitumour activity of an anti-ganglioside GM3(Neu5Gc) antibody. Gene Ther. 2015;22(12):960–7.

61. van den Berg FT, Makoah NA, Ali SA, et al. AAV-Mediated Expression of Broadly Neutralizing and Vaccine-like Antibodies Targeting the HIV-1 Envelope V2 Region. Mol Ther Methods Clin Dev. 2019;14:100–12.

62. Marino M, Holt MG. AAV Vector-Mediated Antibody Delivery (A-MAD) in the Central Nervous System. Front Neurol. 2022;13:870799.

63. Mendiratta G, Jones MK, Stites EC. How often is each gene mutated within the cancer patient population? Mol Cell Oncol. 2022;9(1):2065176.

64. Dromain C, Pavel ME, Ruszniewski P, et al. Tumor growth rate as a metric of progression, response, and prognosis in pancreatic and intestinal neuroendocrine tumors. BMC Cancer. 2019;19(1):66.

65. Ackerman M, Levary D, Tobon G, et al. Highly avid magnetic bead capture: an efficient selection method for de novo protein engineering utilizing yeast surface display. Biotechnol Prog. 2009;25(3):774–83.

66. Boder ET, Wittrup KD. Yeast surface display for screening combinatorial polypeptide libraries. Nat Biotechnol. 1997;15(6):553–7.

67. Chao G, Lau WL, Hackel BJ, et al. Isolating and engineering human antibodies using yeast surface display. Nat Protoc. 2006;1(2):755–68.

68. Fromant M, Blanquet S, Plateau P. Direct random mutagenesis of gene-sized DNA fragments using polymerase chain reaction. Anal Biochem. 1995;224(1):347–53.

69. Imura Y, Tagawa T, Miyamoto Y, et al. Washing with alkaline solutions in protein A purification improves physicochemical properties of monoclonal antibodies. Sci Rep. 2021;11(1):1827.

